# Integration of past caspase activity biases cell elimination *in vivo*

**DOI:** 10.1101/2025.05.18.654727

**Authors:** Tom Cumming, Alexis Villars, Anđela Davidović, Florence Levillayer, Léo Valon, Romain Levayer

## Abstract

The fine tuning of apoptosis in epithelia is essential for regulating tissue size, shape, homeostasis and the maintenance of sealing properties. Regulation of cell death is mostly orchestrated by the activation of Caspases, proteases which were long thought to trigger an irreversible engagement in cell death. However, recent data *in vivo* and *in vitro* outline numerous non-apoptotic functions of caspases as well as quite ubiquitous sublethal activation of effector caspases during development. Yet, it remains unclear in many instances what drives the bifurcation between cell death engagement and cell survival upon caspase activation. The existence of a caspase activity threshold was generally considered to underpin this binary decision, but this was never assessed quantitatively *in vivo* especially at the single cell level. Using quantitative live imaging combined with machine learning and optogenetics in the *Drosophila* pupal notum (a single layer epithelium), we reveal for the first time the existence of a large heterogeneity of caspase sensitivity between cells, as well as the existence of distinct spatial domains with low or high sensitivity to caspases. Using correlative and perturbative experiments, we outline the central role of past exposure to sublethal caspase activity which sensitises cells for apoptosis for several hours. Integrating information about past caspase activation is sufficient to explain most of the global pattern of caspase sensitivity and predict at the single cell level which cells will engage in apoptosis. Finally, we demonstrate that past sublethal caspase activation in a subset of cells is sufficient to bias cell elimination at the clonal and single cell level, thus revealing an alternative mechanism of physiological cell competition. Altogether, this work reveals for the first time the existence of a new layer of apoptosis regulation *in vivo* downstream of effector caspases which can be developmentally regulated and bias clonal selection and the spatial pattern of cell death.

## Introduction

Apoptosis, the most prevalent form of programmed cell death, is essential to modulate tissue shape and size and contributes to development robustness^1, 2^. This is particularly relevant in epithelia, which form a tight mechano-chemical barrier through cell-cell adhesion, and can undergo massive remodelling during development or adult organ homeostasis^3^. While the molecular regulators of apoptosis are very well known, and all converge upon the cleavage and activation of the caspase proteases and the subsequent cleavage of thousands of proteins^4^, we still have a relatively limited understanding of apoptosis regulation in a multicellular and tissular context. This is epitomised by our inability to predict precisely when, where, how many and which specific cells will die in an epithelial tissue, most likely revealing the complex multilayered regulation of epithelial cell death^5^. Nonetheless, this level of predictivity appears essential when it comes to understanding the evolution of tissue size and shape, or to anticipating clonal evolution during development or tumour progression^6^.

The activation of the most terminal caspases, so called effector/executioner caspases, was long though to constitute an irreversible point of engagement in cell death^7-12^. As a result, most molecular dissections of apoptosis regulation have focused on identifying factors acting upstream of caspase cleavage. However, studies over the past decade have now characterised an ever-increasing number of unexpected effector caspase functions which are unrelated to programmed cell death^13-15^. Alternatively, quantitative studies in cell culture revealed early on the variability of cell response to cytotoxic stress, so called fractional killing^16^, where heterogeneity and noise related to cell protein content can alter the engagement in apoptosis despite similar upstream pro-apoptotic cues^17-19^. Similarly, mitochondrial outer membrane permeabilisation (MOMP), which was long thought to be an all-or-nothing process systematically leading to apoptosis^7, 20^, was recently shown to be more nuanced due to the occurrence of partial MOMP that does not lead to cell death^21, 22^. The development of novel genetic sensors of effector caspase activation revealed quite systematic exposure of cells to sub-lethal effector caspase activity *in vivo* in *Drosophila*^23, 24^ and in cancer cell lines^25, 26^. These works clearly question the previously dominant view of caspase activation as the terminal point of engagement in apoptosis, and suggest that a decision process takes place downstream of effector caspase activation. Several alternative models have been proposed to redirect caspase to alternative functions, including the sequestration of active caspases to subcellular compartments^27, 28^, or more generally the existence of a well-defined threshold of activity below which the cell will not enter apoptosis^29^. Accordingly, genetic evidence in *Drosophila* using a combination of caspase mutants and irradiation suggested that the susceptibility to engage in apoptosis could be set by procaspase levels^29^. However, more recent data combining single cell quantification of caspase activity using a live sensor and optogenetic control of caspase found little correlation between caspase activity and the probability to enter apoptosis in HeLa cells^30^. Thus, what determines the decision between survival and death following effector caspase activation remains largely unclear, especially *in vivo*.

The *Drosophila* pupal notum, a single layer epithelium, is an ideal system to study the quantitative regulation of apoptosis in epithelia *in vivo*. During metamorphosis, close to a thousand cells die in less than 24 hours in a highly stereotyped global pattern^31-36^, which can be visualised through long term live imaging while combined with genetic modulation (**Fig. 1A**). Effector caspase activation systematically precedes and is required for cell extrusion toward the basal side^32, 34-38^, which is in turn systematically followed by DNA compaction and apoptosis^36^. Using various live sensors of effector caspases, we previously outlined the large proportion of cells undergoing transient effector caspase activation without committing to cell extrusion and apoptosis^32, 34, 35^, which is also related to the reversion of caspase activity triggered by local pulses of EGFR/ERK activation near extruding cells^35^.

**Figure 1:**
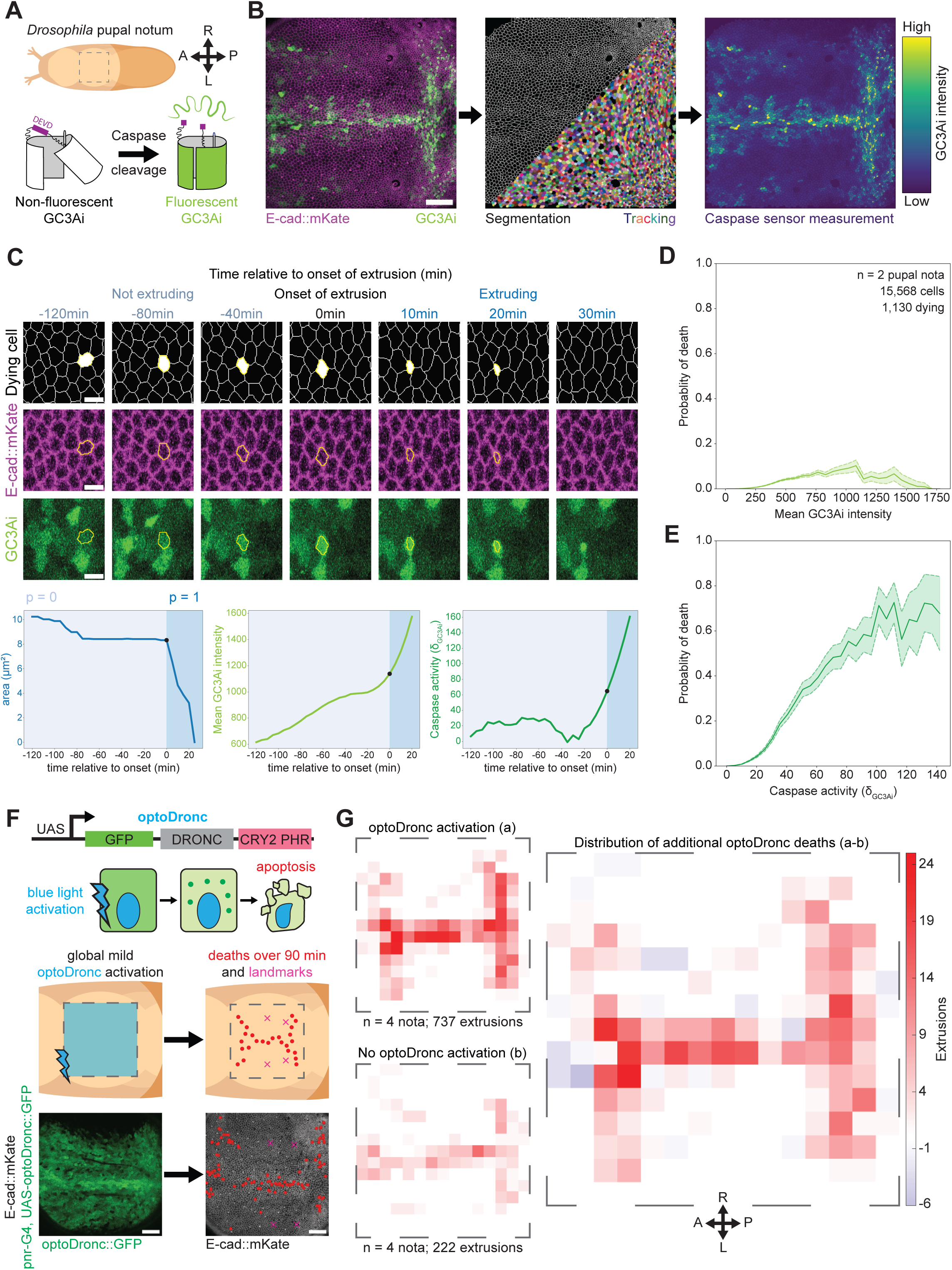
Cell-to-cell and spatial heterogeneity of caspase threshold associated with cell death (associated with Supplementary Figure 1 and 2). **A:** Schematic of the *Drosophila* pupal notum (dotted square) and the GC3Ai effector caspase sensor, which folds and emits fluorescence upon cleavage of the DEVD consensus cleavage site. A: Anterior, P: Posterior, L: Left, R: Right. **B:** Local z-projection of a live pupal notum expressing E-cad::mKate (magenta) and GC3Ai (green) in the pnr domain. Middle panel shows the cell contour upon segmentation with EPySeg, curation with EpiCure, and cell tracking with trackpy. Right panel shows GC3Ai intensity measurement in single cells (blue: low, yellow: high). Scale bar=50μm. **C:** Snapshots of a dying cell before (left, negative time) and after the onset of extrusion (right, positive time), E-cad::mKate in magenta and GC3Ai in green. Yellow polygon shows the cell contour. Scale bars=5μm. Bottom graphs show the corresponding apical area (left), GC3Ai intensity (middle) and estimated caspase activity (right) based on the GC3Ai intensity derivative. The dark blue zone corresponds to the time where the cell is engaged in extrusion, after the area inflection point shown by the black dots. **D,E:** Estimation of the probability to be engaged in cell death (corresponding with measurements taken where the cell is engaged in extrusion, see dark blue area in the curves in **C**) as a function of the current GC3Ai intensity (**D**), or current effector caspase activity, estimated with the GC3Ai intensity derivative (**E**). n= 2 fully segmented pupal nota, 15,568 cells and 1,130 dying cells. While there is a high proportion of non-dying cells at high GC3Ai, there is a good correlation between probability to die and levels of GC3Ai derivative. However, the curve does not correspond to a single threshold value and outlines a large range of values at which cells engage in apoptosis. **F:** Schematic of the opto-Dronc construct, an optogenetic Caspase9 allowing tunable control of caspase activity in *Drosophila*^35^. The *Drosophila* Caspase 9 (Dronc) is fused to GFP in N-ter and CRY2-PHD in C-ter which oligomerises upon blue light exposure (green dots, middle scheme), triggering Dronc cleavage and caspase activation. Bottom scheme illustrates the zone of mild activation of optoDronc used to analyse the spatial pattern of cell death (red dots). The consensus maps of resulting deaths are obtained by Procrustes alignment of several tissues using the aDC and pDC macrochaetae as landmarks (pink crosses). Bottom left shows the expression pattern of optoDronc in the pnr-Gal4 domain and the bottom right picture shows one example of cell death distribution upon optoDronc activation (red dots, cell contour shown with E-cad::mKate, grey). Scale bars=50μm. **G:** Summed map of the extrusions counted in nota for 90 minutes following mild optoDronc activation with blue light pulses, n=4 nota aligned and scaled, 737 extrusions. Bottom map shows the extrusion in control nota (same genotype, no blue pulse, 222 extrusions). The estimation of the distribution of cell death triggered by optoDronc (right map) is obtained by subtracting the control death map (b) from the blue light-exposed map (a). The total local number of deaths is colour coded (red high, blue low). A: Anterior, P: Posterior, L: Left, R: Right.

Here, we use the pupal notum to characterise for the first time quantitatively *in vivo* the relationship between effector caspase activity and commitment to apoptosis. Using a live sensor of effector caspases and a large dataset of tracked cells, we first confirmed that caspase activity provides good predictive power regarding the likelihood of cell death using machine learning (time series forest). However, we surprisingly found that cells commit to apoptosis at various levels of effector caspase activity. Using optogenetic sublethal activation of caspases, we also outlined the reproducible global tissue pattern of caspase sensitivity in the notum, thus confirming that commitment to apoptosis downstream of caspase can be developmentally regulated. Using correlative analysis combined with quantitative optogenetic perturbations, we revealed for the first time that past exposure to caspase can prime cells for apoptosis a few hours later. This contribution of past exposure can explain a significant proportion of the global spatial heterogeneity of caspase sensitivity and is sufficient to predict which cells will commit to apoptosis following a pulse of caspase activation. Finally, using machine learning and clonal perturbation, we confirm that past exposure to caspases can bias cell elimination relative to neighbouring cells and modulate clonal selection during development, hence describing an original mode of physiological cell competition.

## Results

### Spatial and cell-cell heterogeneity of caspase thresholds

We first characterised the effector caspase activity levels associated with engagement in apoptosis in the pupal notum. We segmented and tracked more than 15,000 cells using deep learning pipelines^39^ and a new plugin for segmentation curation and tracking^40^ to measure the temporal dynamics of caspase activity at the single cell level using the GFP-based effector caspase sensor GC3Ai^41, 42^ (**Fig. 1A,B, movie S1**). Given the stability of activated GC3Ai^42^, we used the intensity derivative (the rate of production of active GC3Ai) as a proxy for effector caspase activity^35^ (called “caspase activity” hereafter). Engagement in cell extrusion is systematically followed by DNA compaction and apoptosis^36^. We therefore used the initiation of extrusion as a proxy for engagement in apoptosis and systematically detected the onset of area reduction in extruding cells to define cell death engagement (see Methods and ^37^, **Fig. 1C, movie S2**). We first used machine learning to identify relevant single cell parameters predictive of engagement in apoptosis. To avoid bias coming from unbalanced data and the impact of global tissue patterning on cell shape and behaviour, we trained the models systematically using one dying cell and one of its direct surviving neighbours as a control. To integrate temporal features (mean, standard deviation, derivative) we used a time series forest classifier (TSF^43^, see **Methods**) on backward temporal windows of one hour starting at least 10 minutes prior to the onset of extrusion. We first confirmed that cell apical area and perimeter could provide a good prediction of cell death engagement prior to the onset of extrusion (**Fig. S1 A,C,D**), as previously demonstrated^36^. While cell eccentricity had little predictive value (**Fig. S1B**), we found however that caspase activity could accurately predict cell death (**Fig. S1E,F**), especially in the hour preceding the initiation of extrusion. This suggested that instantaneous caspase activity is relevant to predict the engagement in apoptosis and that the decision to die or survive is taken close to the onset of cell extrusion. We therefore explored more extensively the quantitative behaviour of caspase activity close to the cell death engagement point.

We systematically measured the levels of caspase activity before and after the onset of cell extrusion in dying cells, as well as measuring caspase activity in non-dying cells. Doing so, we obtained the probability of being in apoptosis for a given level of cumulative caspase activity (absolute GC3Ai intensity) or instantaneous caspase activity (derivative of GC3Ai signal, **Fig 1D,E**). We first found that absolute GC3Ai signal on its own provides little predictive information on the engagement in cell death, as exemplified by the large proportion of living cells with high intensity levels (**Fig. 1D, Fig. S1 G,H**, logistic fit Tjur’s R^2^=0.016) and as previously reported in HeLa cells^30^. We observed however a good logistic fit when plotting the proportion of cells engaged in apoptosis as a function of caspase activity (**Fig. 1E, Fig. S1 G,I** logistic fit Tjur’s R^2^=0.301). While this result suggests that effector caspase activity is partially predictive of cell death engagement, it also outlines the significant cell-to-cell variability in the caspase threshold associated with commitment to apoptosis, as the range of caspase activity levels at which cells engage in apoptosis spans nearly two orders of magnitude (**Fig. 1E**). We confirmed that the variability of caspase threshold does not correlate with cell-to-cell heterogeneity of Gal4 expression levels as there was no correlation between caspase at the onset of extrusion and level of expression of another constitutively expressed UAS-driven fluorescent sensor (**Fig S1J,K,** Spearman correlation r= - 0.124, p=0.58).

We then tested whether this cell-to-cell heterogeneity in death-inducing caspase threshold had any developmental relevance. To assess the variability of caspase sensitivity in space, we used an optogenetic initiator caspase previously developed by our group (optoDronc^35^) to trigger mild and uniform activation of caspases in a large band encompassing most of the notum (using the pnr-Gal4 driver, **Fig. 1F**). We then systematically tracked the occurrence of extrusion in the domain expressing optoDronc, and registered the position of macrochaetae (large bristles) from each mapped notum as spatial landmarks for a procrustes transformation, allowing us to scale and align death events from several movies to a single spatial map (see **Methods** and **Fig. S2A**). The same procedure was applied to pupae with the same genotype in the absence of blue light pulses as controls. We first revealed that the vast majority of additional deaths occurs in the 90 minutes following mild activation of optoDronc (**Fig. S2B, movie S3**). We therefore created maps of cell death using extrusions occurring in the first 90 minutes following blue light pulses. We subtracted the distribution of spontaneous death in the control map from the blue light-stimulated map to obtain the spatial distribution of additional deaths triggered by the activation of optoDronc (**Fig. 1G, movie S3**). This revealed a clear heterogenous pattern of death induction with a higher susceptibility to caspase in the midline and in bands at the anterior and posterior edges of the tissue (**Fig. 1G, movie S3**). This heterogeneity is not explained by spatial variability of the driver (**Fig 1F bottom**), nor by differences in the kinetics of effector caspase activity downstream of optoDronc (no spatial difference of GC3Ai dynamics upon prolonged blue light activation, **Fig S2C, movie S4**). Moreover, the comparison of the cumulative number of deaths over 3 hours in the controls and the nota exposed to blue light clearly shows that optoDronc pulses trigger a net increase in the number of cell deaths (**Fig. S2B,** 956 versus 482 extrusions, 4 pupae each). This confirmed that the pattern we observed are “additional” deaths events and not a simple “acceleration” of the elimination of cells that would have died during normal development. Thus, we revealed a stereotypical spatial pattern of caspase sensitivity, which suggests that the engagement in apoptosis downstream of effector caspases can be developmentally regulated.

### Past caspase activity primes cells for apoptosis

We next set out to identify parameters that could explain the heterogeneity of caspase sensitivity. We isolated the 1,130 dying cells from our dataset, extracted the level of caspase activity at the onset of extrusion of each cell, and assessed the strength of the correlation between this value and various morphometric and caspase-related parameters in windows of time prior to extrusion (**Fig. 2A,B**). Doing so, we outlined an interesting negative correlation between GC3Ai intensity in the past (up to 4-5 hours prior to extrusion) and caspase activity at the onset of extrusion (**Fig. 2C**). This suggested that previous exposure to caspase activity may decrease the instantaneous caspase threshold required to enter apoptosis, hence priming the cell for death. We therefore experimentally tested whether previous exposure to sublethal caspase could indeed prime cells for apoptosis. We used optoDronc to locally expose a subdomain of the notum (lateral to the midline where there is no physiological cell death^33, 35, 36^) to two pulses of caspase stimulation separated by 1, 3 or 5 hours, and used as a control the contralateral region exposed only to the latter stimulation (**Fig. 2D, movie S5**). We observed that cells were significantly more likely to die in the region with prior caspase exposure compared to the single exposure region, with a more than 4-fold increase in cell death number after 1h, and ∼25% increase after 3 hours (**Fig. 2E, movie S5**). This confirmed that past exposure to caspase can indeed prime cells for apoptosis for a few hours.

**Figure 2:**
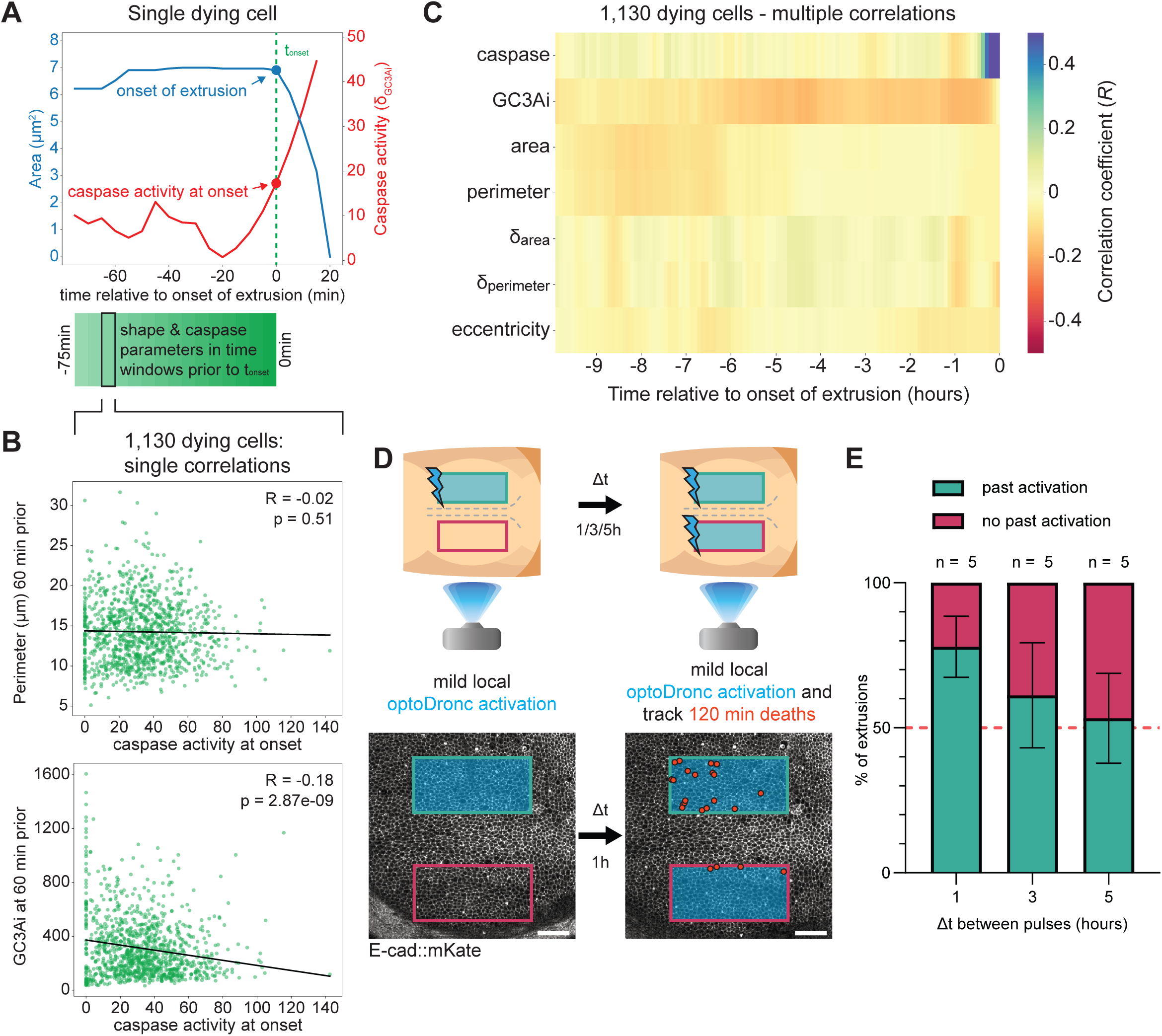
Past caspase activation primes cells for apoptosis. **A:** Cell area (blue) and caspase activity (GC3Ai intensity derivative, red) of a single dying cell. The green dotted line shows the onset of cell extrusion. The blue and red dots show the area and caspase activity at the onset of apoptosis engagement. We tracked multiple parameters backward in time (green squares, 20 minutes sliding window shifted every 5 minutes) related to cell geometry and caspase dynamics on 1,130 dying cells (2 nota). **B:** Examples of correlations calculated between caspase activity at the onset of apoptosis engagement and perimeter from 70 to 50 min prior to apoptosis engagement (top) and GC3Ai intensity from 70 to 50 min prior to apoptosis engagement (bottom). **C:** Matrix of correlations of various geometrical (cell apical area, perimeter, rate of change of area, rate of change of perimeter, eccentricity) and caspase-related parameters (GC3Ai intensity, caspase=derivative of GC3Ai intensity) with caspase activity at the onset of engagement in apoptosis. Correlations are calculated on time windows of 20 minutes prior to the onset of cell extrusion (t=0 hours). Blue: positive correlation, Red: negative correlation. Note the clear global negative correlation between GC3Ai intensity in the past and caspase at the onset. n=1,130 dying cells, 2 nota. **D:** Schematic of the experiment directly testing the influence of past caspase activation on apoptosis engagement. *pnr-Gal4* drives *UAS-optoDronc* which is activated with a short blue light pulse in the lateral right part of the notum (top, regions with very limited spontaneous cell death). After different waiting times (Δt; 1, 3 or 5 hours), the left and right regions of the notum are exposed to blue light. The number of extrusions is then compared in the left and right region for the next 120 min. Bottom pictures show an example of a notum with the distribution of cell extrusions in the left and right regions over 120min following the second blue light pulse (one red dot=one cell extrusion) Scale bars=50μm. **E:** Quantification of the proportion of cell extrusions in the right (twice-exposed) and left (once-exposed) regions, as a percentage of the total extrusions per notum in the 120 minutes following the second blue light pulse, with a waiting time of 1, 3 and 5 hours after the first blue light exposure (n = 5 nota per condition). Error bars show s.e.m..

### Integration of past caspase activity is predictive of the spatial pattern of death and single cell engagement in apoptosis

So far our results suggested that past caspase activity provides relevant information to predict the engagement in apoptosis upon further caspase activation. We therefore revisited our experiments triggering global mild caspase activation in the notum (**Fig. 1F,G**), and noticed that the pattern of apoptosis induction closely matched the pattern of absolute GC3Ai intensity (compare **Movie S1**, **Fig. 1B** with **Fig. 1F**). We therefore checked whether the pattern of cumulative past caspase activity (using absolute GC3Ai intensity as a proxy) could indeed explain the preferential localisation of cell death following a pulse of optoDronc activation. The pattern of GC3Ai intensity changes during pupal development, with early pupae showing a strong signal in the anterior domain, mild signal in the midline and no signal in the posterior region (∼18 hours APF), while at late stage there is a strong increase in signal in the midline and the posterior region (∼24 hours APF, **movie S1, Fig. 3A**). We therefore used optoDronc to trigger mild global caspase activation in early and late pupae to compare the pattern of cell elimination and the pattern of GC3Ai intensity. Strikingly, caspase activation in the early pupae mostly triggers death in the anterior and midline domain and has little impact in the posterior region, while induction of caspase in late pupae triggered death in the anterior, midline and posterior regions, correlating well with the corresponding pattern of GC3Ai signals at the relevant developmental stage (**Fig. 3A**, **movie S6** Pearson correlation comparing the spatial pattern of GC3Ai intensity and local extrusion number r=0.31, p=1.60.10^-6^, for early pupae and r=0.65, p=1.67.10^-28^ for late pupae). This suggested that the pattern of death induction by optoDronc may be largely explained by past exposure to caspase activation. To directly confirm the link between past activation and the spatial pattern of caspase sensitivity, we erased physiological caspase activity by depleting the pro-apoptotic gene *hid* using RNAi^34, 37^ (see the almost absence of GC3Ai signal **Fig. 3B** left, **movie S7**), and triggered a pulse of caspase activity in the *pnr* domain with optoDronc (**Fig. 3B**, **movie S6**). Epistatically, optoDronc can directly trigger caspase activation downstream of the pro-apoptotic genes *hid*, *grim* or *reaper^5^* and bypass the depletion of *hid* by RNAi. We therefore characterised the pattern of death induction by optoDronc in the *hid* RNAi background. Interestingly, the pattern of cell elimination was much more homogeneous in the context of *hid* RNAi compared to optoDronc pulses in WT background (**Fig. 3B**, compare to **Fig 3A**). The diminution of spatial patterning is reflected by the considerable reduction in clustering of cell death distribution in the *hid* RNAi context compared to late controls (**Fig. 3C**, reduced global Moran’*I*). This confirmed that past caspase activation (here regulated by *hid*) is essential to generate the spatial bias of cell elimination. We noticed however that there is still a slight increase in the probability to observe cell death near the midline and in the anterior domain, accordingly the spatial autocorrelation is still higher than that expected from a random spatial distribution (**Fig. 3C**, global Moran’s *I* for a random distribution ≈ 0). This suggests that other unknown spatially patterned factors can also modulate the susceptibility to caspases in the notum independently of past caspase activity.

**Figure 3:**
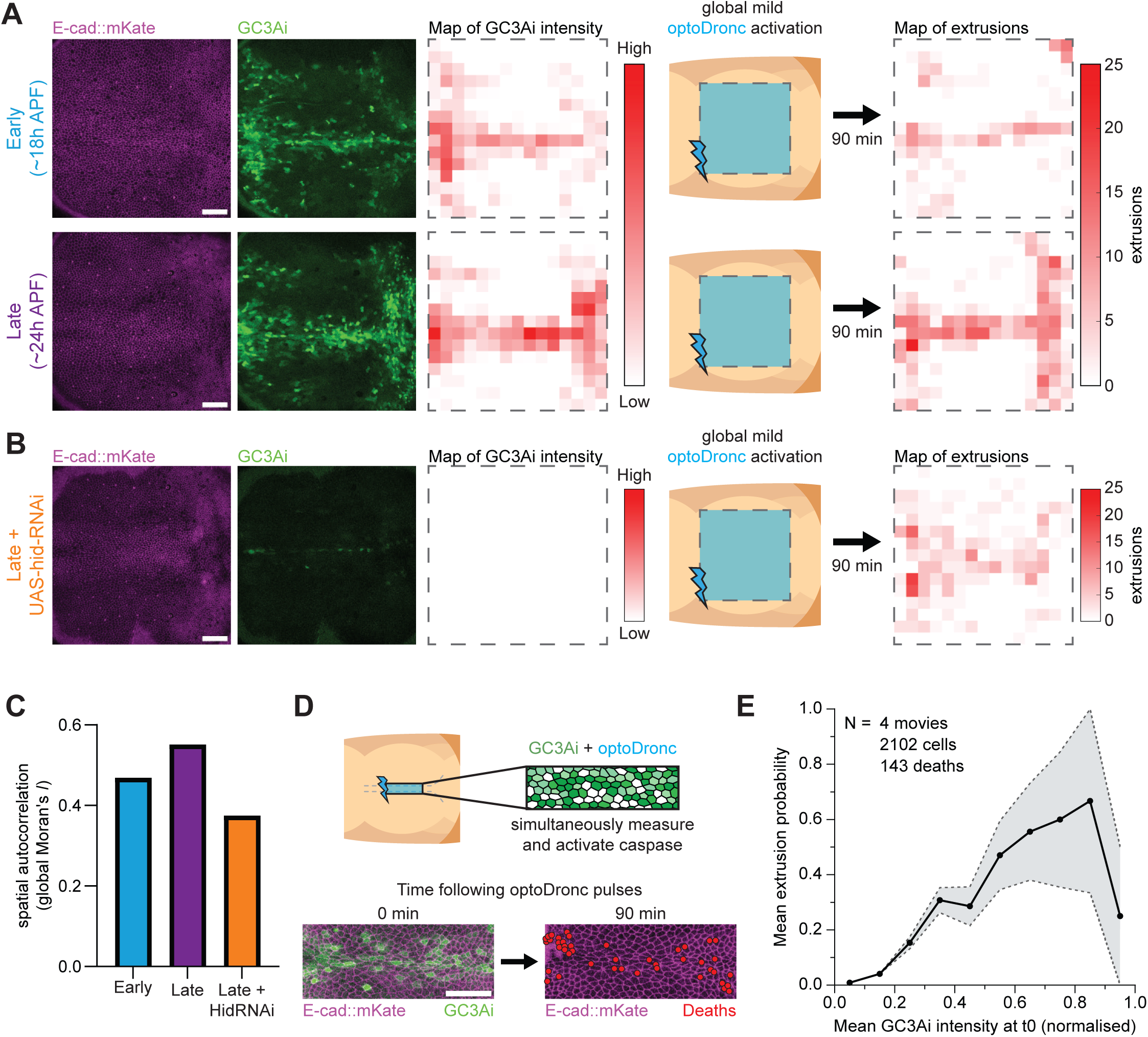
Past caspase activation is predictive of apoptosis spatial pattern and single cell decision. **A,B:** Left, local projections of an early (top, ∼18h APF), and late (middle, ∼24h APF) pupal notum (**A**), or late pupal notum upon depletion of the pro-apoptotic gene *hid* by RNAi (bottom, **B**). E-cad::mKate (magenta), *pnr-Gal4 UAS-GC3Ai* (green, same intensity scale for all) and the spatial map of the GC3Ai intensity (right, white to dark red, same scale in the three conditions). Scale bars=50μm. Middle: schematic of the global optoDronc activation performed and the tracking of cell death over 90 minutes. Right: Sum maps of cell death distribution observed in the 90min following blue light pulse obtained after the rescaling/alignment of 3 young (top, **A**), 3 old (middle, **A**), and 3 *UAS-hid dsRNA* pupae (bottom, **B**), total extrusion number: 238 in the young, 644 in the old pupae, 345 in old *UAS-hid dsRNA* pupae. Colour scale (white to dark red, low to high number of extrusions) is the same in the 3 conditions. **C:** Spatial autocorrelation of the extrusion pattern observed over 90min after optoDronc activation in young, old, or *UAS-hid dsRNA* pupae (see **Methods**). **D:** Top, schematic of the experiment, local activation of optoDronc in the midline is combined with the imaging of GC3Ai intensity at t0. Cell extrusion is then mapped in the next 90min. Bottom, local projection of an early pupa at the onset of blue light exposure (GC3Ai signal, green, E-cad::mKate, magenta). The localisation of cell death in the next 90min is shown with red dots. Scale bar=50μm. **E:** Single cell probability of cell extrusion during the 90min following optoDronc activation in the midline as a function of the normalised GC3Ai raw intensity at t0. Grey area is the standard error of the mean (note that probability measurement is poorly reliable above 0.7 due to the low number of cells showing these extreme values).

We then checked whether the history of cells and their previous exposure to caspase could also be predictive of the engagement in apoptosis at the single cell level. Combining both optoDronc and GC3Ai under the expression of the notum-wide pnr driver, we used a pulse of blue light to simultaneously image the levels of GC3Ai at time 0 and globally activate caspase. We then assessed which cells will die in the midline, and found a striking correlation at the single cell level between the probability to die and the levels of GC3Ai intensity at time 0 (**Fig. 3D,E, movie S8**). This suggests that cumulative past exposure to caspase is predictive of which cell will engage in apoptosis following a pulse of caspase activity. While we previously showed that absolute GC3Ai levels on their own are poorly predictive of apoptosis engagement (**Fig. 1D**), we show here that cumulative past caspase activation (GC3Ai levels) can significantly modulate cell elimination probability when caspase becomes active at a given time. Altogether, these results suggest that the decision to die is first dominated by the current status of caspase activity, which is then fine-tuned by cumulative past exposure.

### Relative past and current caspase activity can bias clonal elimination and the death probability relative to neighbours

We previously showed that the initiation of cell extrusion triggers a pulse of ERK activity in the neighbouring cells which shuts down caspase and protects them from apoptosis^35^. Accordingly, when neighbouring cells activate caspase simultaneously, the first cell to enter extrusion will inhibit the death of its neighbours^35^ (**Fig 4A**,), in a similar fashion to cell selection by lateral inhibition^44^. As a result, cell death decision-making is not fully cell autonomous and rather relies on which cells in the group will be the fastest to enter extrusion/apoptosis. We therefore tested whether relative differences in caspase activity and past exposure to caspase could bias the selection of the cell to die among neighbours. We compared dying cells and their neighbours and checked which parameter is predictive of the first cell to die. Among 1,130 groups of individual dying cells and their neighbours, the first cell to die had the highest GC3Ai signal in 52% of cases (random assignment would be close to 15%) and had the highest caspase activity in 77% of cases (**Fig 4B**, comparison at the onset of the first extrusion). This suggested that relative caspase activity and relative previous exposure to caspase could both bias cell elimination among neighbours. To confirm that our capacity to predict the engagement in death among a group of cells is not just restricted to the timepoint at the onset of extrusion, we again used TSF machine learning to test the predictive strength of relative differences in caspase between neighbouring cells over time. We first used the TSF model to assign a death probability value for each cell in a group of neighbours, and assumed that the one with the relative higher probability score should be the one to die first (**Fig. 4C**. see **Methods**). We used the relative GC3Ai and caspase activity values between the dying cell and its neighbours for training the TSF model. This led to very good prediction of which cell will die first in a group when using the first hour window preceding cell extrusion onset with GC3Ai intensity or caspase activity (**Fig. 4C**, F1 scores>0.7, note that TSF of GC3Ai also includes information about the derivative), and stronger predictive capacity for raw GC3Ai starting 1 hour prior to the onset of extrusion and lasting up to 3 hours (**Fig. 4C**). Altogether, these results confirmed that the combination of the relative values of caspase activity and past exposure are sufficient to predict which cell will die first in a group of neighbours.

**Figure 4:**
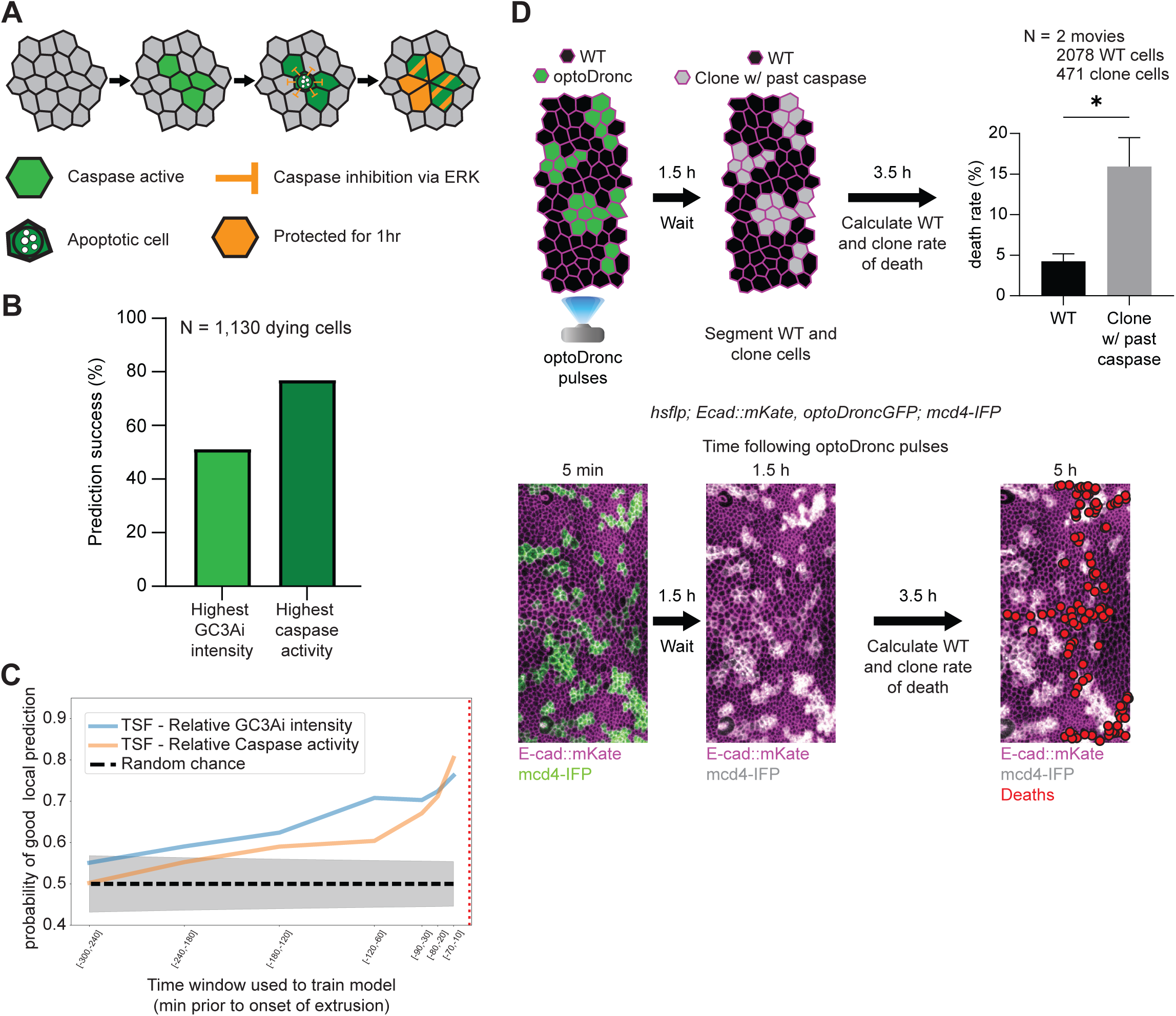
Past caspase activation biases single and clonal cell elimination. **A:** Schematic of local ERK negative feedback on epithelial cell death. Upon initiation of cell extrusion, the activation of ERK in the neighbouring cells (orange) blocks caspase activity (green) and protect them from cell death for at least 1 hour (see ^35^). **B:** Percentage of prediction success regarding which cell will die first among a group of neighbouring cells based on comparison of GC3Ai/caspase parameters at the onset of the first extrusion event. The first dying cell has the highest GC3Ai signal in 52% of cases, and the highest caspase activity in 77% of cases. n=1,130 groups of cells. **C:** Proportion of correct prediction of the first cell dying among a group of neighbours based on a Time Series Forest model. The cell with the highest probability score in the group is predicted to be the first to die. TSF is trained either on relative GC3Ai intensity (blue) or on relative caspase activity (orange) comparing a dying cell and a surviving neighbour as control (see **Methods**). The black dotted line shows the expected score for random prediction, the grey area shows the 95% confidence interval (prediction is tested on a balanced dataset of dying cells and surviving neighbours, a random prediction will be correct in 50% of cases). The time windows (starting from 10 minutes prior to extrusion onset) used for TSF prediction and training are shown on the x-axis (time in minutes prior to extrusion onset). **D:** Schematic of the experiment triggering sublethal caspase activation in clones. Clones expressing optoDronc (green) located in the posterior region of the pupal notum are exposed to mild blue light pulses. Rate of death is then calculated in the clonal cells and outside the clones from 1.5hrs to 5hrs after blue light pulses. In this time window, cell death is mostly triggered by spontaneous developmental activation of caspase in the posterior region. Bottom panels show local projection of a pupal notum expressing E-cad::mKate (magenta) and UAS-mCD4::mIFP (green/grey) at the onset of blue light pulses, 1.5hrs and 5hrs later. Red dots show cell extrusion events counted from 1.5hrs to 5hrs. The death probability in control and optoDronc clones is shown in the bar graph (n=471 clonal and 2078 WT cells, 2 nota, p=5.31e^-17^, Fisher’s exact test). Error bars are 95% confidence interval. Scale bars=30μm.

This non-cell autonomous death decision is reminiscent of cell competition^45, 46^, a context-dependent cell elimination process which triggers the elimination of suboptimal cells by their neighbours through short range communication. We therefore checked whether past exposure to caspase could indeed bias clonal selection during normal development. We triggered mild/sublethal activation of optoDronc in clones of the posterior domain of the notum and quantified the probability of cell death from 90min to 5 hours after blue light pulses (hence not counting deaths directly triggered by the first pulse of activation, see **Fig S2B**). We found a striking increase in death probability relative to neighbouring WT cells (**Fig 4D, movie S9**, 3.5-fold increase of cell death probability), confirming that past exposure to caspase is sufficient to generate “loser”-like clones which are more likely to be eliminated in regions where caspases become physiologically activated later. Altogether, we showed that past exposure to caspase can bias single cell elimination and clonal dynamics, suggesting that the history of caspase activation could be sufficient to generate physiological cell competition by priming a subset of cells for apoptosis. This novel mechanism of physiological cell competition would not require direct comparison between cells and could be sufficient to bias cell elimination.

## Discussion

We outlined here for the first time the large diversity and spatial heterogeneity of caspase sensitivity *in vivo*. The existence of reproducible large spatial patterns of caspase sensitivity suggests for the first time that the decision to engage in apoptosis downstream of effector caspases can be developmentally regulated. Using machine learning, optogenetics and correlative analysis, we demonstrated that past exposure to caspase activation can prime cells for apoptosis in the following hours. This “memory” of past activation explains a significant proportion of the spatial bias of caspase sensitivity and can predict which cell will engage in apoptosis following a pulse of caspase activation. Globally, our work demonstrates that while instantaneous caspase activity is the primary predictor of cell death engagement (**Fig 1E** and **Fig. S1**), information provided by past exposure generates a significant proportion of the heterogeneity that can bias cell elimination. Of note, we still found a significant spatial bias of cell death distribution upon optoDronc activation despite erasure of past caspase activity (**Fig. 3B,C**). This suggests that other parameters, yet to be identified, can also modulate the susceptibility to enter apoptosis following caspase activation in the notum.

Previously, quantitative characterisation of caspase dynamics in HeLa cells using the same GC3Ai sensor suggested that caspase activity was poorly predictive of apoptosis engagement^30^, in contrast with our system where the combination of present and past caspase activity is sufficient to predict cell death. This apparent discrepancy may be explained by various differences between the biological systems and the assays. For instance, we focus here on epithelial cells where cell death decision is intrinsically connected to cell extrusion. Moreover, while we can use the engagement in extrusion to temporally align dying cells, there is no trivial criteria to define the onset of apoptosis in HeLa cells and align the temporal profiles, which may have masked relevant information about the utility of caspase activity to predict cell death. Since our machine learning approach suggests that the information provided by caspase activity is mostly relevant in the few hours preceding cell extrusion, such temporal alignment is therefore essential to identify relevant signatures. Finally, the intrinsic resistance of HeLa cells to apoptosis^47-49^ may mask the instructive role of caspases. Accordingly, caspase signals become predictive in HeLa cells when they are primed by stress conditions^30^. Irrespective of these differences, these two studies clearly outline the variability of cell death decision-making following caspase activation and the existence of downstream parameters that can modulate apoptosis engagement. Of note, the influence of past caspase activity on the spatial pattern of caspase sensitivity does not seem to be a peculiarity of the pupal notum. Similar spatial bias can be observed in the wing imaginal disc upon homogeneous and transient pulses of pro-apoptotic genes, which preferentially trigger cell death in regions which were already physiologically exposed to caspases^50^ (**Fig. S3**). This study relies on the relationship between engagement in extrusion and engagement in apoptosis. Interestingly, we observed a systematic exponential increase in caspase activity as individual cells progress in extrusion (^35^ and **Fig. 1C**), which may be compatible with a positive feedback loop whereby extrusion remodelling promotes caspase activity which then promotes extrusion progression. This amplification loop could explain the binary decision associated with extrusion and apoptosis engagement. Future work will help to identify the factors promoting this rapid increase of caspase activity and its relationship with the well-described process of anoikis, whereby loss of adhesion drives cells to irreversibly commit to die^51^.

What, then, are the molecular mechanisms encoding the memory of past caspase activation, and what sets this characteristic timescale of a few hours? This memory is unlikely to be explained by the perdurance of active caspases which were previously shown to be very short-lived^9, 52-54^, in agreement with our previous measurement of the short time delay between pulses of ERK and inflection of caspase activity (averaged delay of 15min^35^) which is not compatible with long-lived activated caspases. Alternatively, partial cleavage of caspase substrates facilitating the engagement in extrusion may explain this apoptotic priming. In this scenario, the timescale of memory should be related to the characteristic turnover of the targeted proteins. Accordingly, we tested different pathways known to be modulated downstream of caspases that could feed-back on apoptosis susceptibility. For instance, JNK pathway activity can be tuned by caspase^55^ and directly impact apoptosis engagement. However, while a fluorescent reporter of JNK (TRE-RFP^56^) shows activation in the midline, there was absolutely no sign of activation in the posterior domain which is exposed to caspase activity for several hours during development of the notum (**Fig. S4A**, **movie S10**). Alternately, reactive oxygen species (ROS) can be triggered by caspases^28, 38, 57-60^ and could modulate apoptosis by increasing cellular toxicity. However, we could not detect clear activation of ROS in the posterior region using a transcriptional reporter (GstD-GFP, **Fig. S4B**, **movie S11**), nor any induction of ROS following a sublethal pulse of optoDronc in one heminotum (**Fig. S4C**). Proteasome inhibition was also shown to facilitate apoptosis engagement through the stabilisation of active caspases^9^, and several proteasome components were shown to be cleaved by caspases in S2 cells^61^. However, a live sensor of proteasome activity (proteoflux^62^) did not reveal distinctive change of protein degradation rate in the posterior domain (**Fig. S4D, movie S12**). Sublethal caspase activity can also modulate cytoskeleton organisation^13^, hence modulating the probability to enter extrusion. Accordingly, sublethal Dronc activity was shown to downregulate MyoII in the pupal notum^63^, and global tension reduction was recently shown to be permissive for cell extrusion^64^. However, we could not detect clear modulation of MyoII intensity following sublethal pulses of optoDronc (**Fig. S4E, movie S13**). We also previously showed that disassembly of microtubules by caspases promotes cell deformability and the initiation of cell extrusion^37^. However, we could not detect any clear change in microtubule organisation upon sublethal optoDronc activation (**Fig. S4F**, **movie S14**). Finally, we tested whether the pattern of Diap1 (the orthologue of XIAP1, a core inhibitor of apoptosis which can directly tune effector caspase activity^65-67^) could partially explain the spatial variability of optoDronc sensitivity. We designed a knock-in line of Diap1 tagged with fluorescent mScarlet to assess the variation of protein levels (since Diap1 can be regulated by protein degradation^66^). We observed a narrow band in the midline with low Diap1, as previously reported using a reporter of Diap1 degradation^68^, and a decrease in Diap1 levels at the posterior edge of the tissue at late stage of development (**Fig. S4G**, **movie S15**). However, we saw no change in Diap1 levels following a sublethal pulse of optoDronc (**Fig. S4H, movie S16**). Thus, we could not identify a single obvious candidate explaining how past caspase activation sensitises cells for apoptosis, which may be compatible with a subtle and combinatorial contribution of the modulation of many caspase targets.

Interestingly, this work outlined the numerous cells which transiently activate effector caspases and yet survive. It remains unclear at this point whether such transient activations represent a failure to engage in apoptosis, a process akin to anastasis^24, 69, 70^, or whether they may also have a relevant physiological function in regulating cytoskeleton/shape or other cellular properties. What, then, could be the physiological relevance of the heterogeneity of caspase sensitivity? At the population level, the existence of heterogeneity and an almost linear correlation between probability to enter extrusion and caspase activity (**Fig. 1E**) suggests that caspase acts like a rheostat rather than a switch at the population level. The heterogeneity of caspase threshold would therefore be physiologically relevant as it would avoid massive simultaneous elimination of cells passing a threshold of caspase activity (hence leading to tissue rupture^35, 71^) and instead progressively eliminate increasing proportions of cells.

Our work also suggests that the engagement in apoptosis is not totally cell-autonomous and relies on which cells among neighbours will engage in extrusion first. In this context, cells that were previously exposed to caspases and/or having relatively higher caspase activity are more likely to die first, either in a physiological context, or upon mild caspase activation in clones. This non-autonomous decision is reminiscent of the process of cell competition whereby neighbouring cells trigger the elimination of suboptimal cells^72^. Interestingly, previous work in mouse embryonic stems cells already suggested that an increased susceptibility to apoptosis is sufficient to generate so-called “loser” cells that will be preferentially eliminated in a developmental context with high apoptosis rate^73, 74^. While *Drosophila* has been a central system to decipher the concepts and mechanisms of cell competition^75^, up till now there has been very little characterisation of spontaneous cell competition taking place during normal development. We propose that the relative modulation of caspase sensitivity by past exposure could constitute one example of such physiological cell competition which could bias clonal selection and the genetic make-up of adult tissues. Future work will help to characterise the contribution of such bias in pre-tumoural clone expansion and cancer progression.

## Acknowledgements

We would like to thank members of the CDEH group for constructive comments on the manuscript. We would also like to thank Anne Loap for her contribution at the start of the project during her internship, Vaishnav Manoj for testing the role of protein stability during his internship, and Georgy Sapozhnikov for initiating the machine learning approach during his internship. We would like to thank Eugenia Piddini for sharing the hs-CL1::GFP line, the Bloomington Drosophila Stock Center and their NIH funding (P40 OD018537), the Drosophila Genetic Resource Center and the Vienna Drosophila Resource Center, Flybase for sharing essential information, stocks and reagents. We thank Gaëlle Letort for developing and sharing the EpiCure pipeline, Alexis Matamoro-Vidal for help with the analysis of apoptosis pattern in the wing discs. We would like to thank the Image Analysis Hub (IAH) of Institut Pasteur for developing the local z-projector plugin and Zellige. TC was supported by a grant from the PPU program of Institut Pasteur and a 4^th^ year PhD grant from the Ligue Nationale Contre le Cancer. Work in RL lab is supported by the Institut Pasteur, the ANR PRC CoECECa, the ANR PRC MAPEFLU, the ANR-10-LABX-0073, the CNRS (UMR 3738) and the European Union (ERC, PrApEDoC, 101085444). Views and opinions expressed are however those of the author(s) only and do not necessarily reflect those of the European Union or the European Research Council. Neither the European Union nor the granting authority can be held responsible for them. A CC-BY public copyright license has been applied by the authors to the present document and will be applied to all subsequent versions up to the Author Accepted Manuscript arising from this submission, in accordance with the grant’s open access conditions.

## Authors contributions

TC, AV and RL designed the project. AV performed the preliminary experiments showing the heterogeneity of caspase threshold and the role of past activation, and participated in large movies segmentation. TC did all the other experiments and analysis. AD performed the statistical analysis of the segmented dataset and the machine learning. FL designed crosses and fly lines. RL and LV supervised the work, RL obtained funding. All the authors participated in the article writing and editing.

## Star Methods

### Resource availability

#### Lead contact

Further information and requests for resources and reagents should be directed to and will be fulfilled by the lead contact, Romain Levayer.

#### Material availability

All the reagents generated in this study will be shared upon request to the lead contact without any restrictions.

#### Data and Code availability

All code generated in this study and the raw data corresponding to each figure panel (including images and local projection of movies) will be accessible on a repository upon final publication of the article. Further information about the dataset can be requested from the lead contact.

### Experimental model and subject details

#### Drosophila melanogaster husbandry

All the experiments were performed with *Drosophila melanogaster* fly lines with regular husbandry techniques. The fly food used contains agar agar (7.6 g/l), saccharose (53 g/l) dry yeast (48 g/l), maize flour (38.4 g/l), propionic acid (3.8 ml/l), Nipagin 10% (23.9 ml/l) all mixed in one litre of distilled water. Flies were raised at 25°C in plastic vials with a 12h/12h dark light cycle at 60% of moisture unless specified in the legends and in the table below (alternatively raised at 18°C or 29°C). Females and males were used without distinction for all the experiments. We did not determine the health/immune status of pupae, adults, embryos and larvae, they were not involved in previous procedures, and they were all drug and test naïve.

#### Drosophila melanogaster strains

The strains used in this study and their origin are listed in the table below.

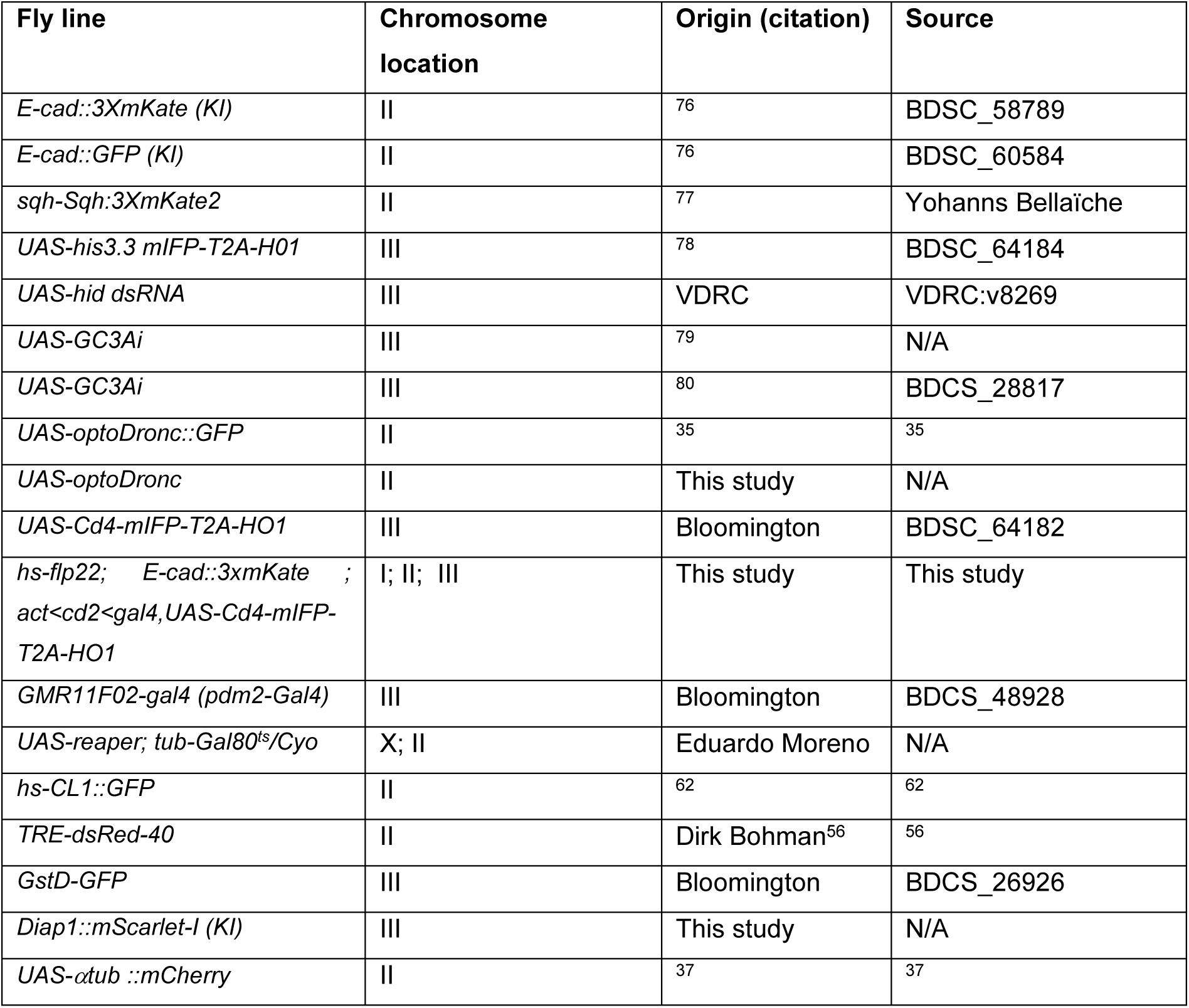

The exact genotype used for each experiment is listed in the next table. ACI: time After Clone Induction, APF: After Pupal Formation, AEL: After Egg Laying.

| Figure | Genotype | Heat shock 37°C | Stage |
| --- | --- | --- | --- |
| 1A-E | *w; E-cad::3xmKate (KI)/UAS-GC3Ai ; pnr-Gal4/+* | N/A | 16h APF |
| 1F,G | *w; E-cad:: 3x mKate (KI)/UAS-optoDronc::GFP ; pnr-Gal4/+* | N/A | 20h APF |
| S1A-H | *w; E-cad:: 3x mKate (KI)/UAS-GC3Ai ; pnr-Gal4/+* | N/A | 16h APF |
| S1J | *w; E-cad:: 3x mKate (KI)/UAS-GC3Ai ; pnr-Gal4/UAS-His3::mIFP-T2A-HO1* | N/A | 16h APF |
| S2A,B | *w; E-cad:: 3x mKate (KI)/UAS-optoDronc::GFP ; pnr-Gal4/+* | N/A | 20h APF |
| S2C,D | *w; E-cad:: 3x mKate (KI)/UAS-optoDronc ; pnr-Gal4/UAS-GC3Ai* | N/A | 20h APF |
| 2A-C | *w; E-cad:: 3x mKate (KI)/UAS-GC3Ai ; pnr-Gal4/+* | N/A | 16h APF |
| 2D,E | *w; E-cad:: 3x mKate (KI)/UAS-optoDronc::GFP ; pnr-Gal4/+* | N/A | 20h APF |
| 3A | *w; E-cad:: 3x mKate (KI)/UAS-GC3Ai ; pnr-Gal4/+* | N/A | 18h and 24h APF |
| 3A,C | *w; E-cad:: 3x mKate (KI)/UAS-optoDronc::GFP ; pnr-Gal4/+* | N/A | 18h and 24h APF |
| 3B | *w; E-cad:: 3x mKate (KI)/UAS-GC3Ai ; pnr-Gal4/UAS-hid dsRNA* | N/A | 18h and 24h APF |
| 3B,C | *w; E-cad:: 3x mKate (KI)/UAS-optoDronc::GFP ; pnr-Gal4/UAS-hid dsRNA* | N/A | 18h and 24h APF |
| 3D,E | *w; E-cad:: 3x mKate (KI)/UAS-optoDronc ; pnr-Gal4/UAS-GC3Ai* | N/A | 20h APF |
| 4B,C | *w; E-cad:: 3x mKate (KI)/UAS-GC3Ai ; pnr-Gal4/+* | N/A | 16h APF |
| 4D | *hs-flp22; E-cad:: 3x mKate (KI)/UAS-optoDronc::GFP ; act<cd2<Gal4, UAS-Cd4::mIFP-T2A-HO1/+* | 48h ACI | 24h APF |
| S3A,B | *UAS-reaper/+ ; tub-Gal80<sup>ts</sup>/+ ; pdm2-Gal4/+* | N/A | 96h AEL |
| S4A | *w; E-cad::GFP (KI)/TRE-dsRed* | N/A | 16h APF |
| S4B | *w; E-cad::GFP (KI)/+; GstD-GFP/+* | N/A | 16h APF |
| S4C | *w; E-cad:: 3x mKate (KI)/UAS-optoDronc; pnr-Gal4, UAS-Cd4::mIFP-T2A-HO1/GstD-GFP* | N/A | 20h APF |
| S4D | *w; E-cad:: 3x mKate (KI)/hs-CL1::GFP* | N/A | 16h APF |
| S4E | *w; sqh:: 3x mKate /UAS-optoDronc; pnr-Gal4, UAS-Cd4::mIFP-T2A-HO1/+* | N/A | 20h APF |
| S4F | *w; UAS-αtub::mCherry/UAS-optoDronc; pnr-Gal4, UAS-Cd4::mIFP-T2A-HO1/+* | N/A | 20h APF |
| S4G | *w; E-cad::GFP (KI)/+; diap1::mScarletl (KI)/+* | N/A | 16h APF |
| S4H | *w; UAS-optoDronc/+; pnr-Gal4, UAS-Cd4::mIFP-T2A-HO1/diap1::mScarletl (KI)* | N/A | 20h APF |

### Generation of fly lines

#### Knock-In Diap1::mScarlet-I

CRSPR-mediated homologous recombination was achieved by injecting three plasmids, one corresponding to the donor template and two encoding the guide RNAs (gRNAs), into Cas9-expressing embryos (BL55821) by BestGene Inc and screened for 3xP3-DsRed insertion. The following gRNAs were selected using the Target Finder tool (https://flycrispr.org/target-finder/). The guide sequences (with PAM sequence underlined) were:

gRNA1, 5′-GGCTCCGTCCGTAGCTCCCACGG -3′

gRNA2, 5′-GGTTGATGAGTAAGCTACATGGG -3 ′.

Corresponding oligonucleotides containing the guide were annealed and cloned into BsbI digested pU6.3-gRNA

(DGRC Stock 1362; https://dgrc.bio.indiana.edu//stock/1362; RRID:DGRC_1362).

To produce the homologous repair construct pScarlessHD-Diap1-mScarlet-DsRed, all cloning was performed by using NEBuilder HiFi DNA Assembly Method. Linear DNA fragments were amplified by PCR and inserted into pScarlessHD-DsRed (DGRC #1364) see https://flycrispr.org/scarless-gene-editing/.

The final construct was checked by sequencing.

The homology arms (1kb long) flanking the target sites were amplified from genomic DNA extracted from line BL55821, the mScarlet-I was amplified from pmScarlet-i_C1 (Addgene 85044). The candidate lines were sequenced to confirm successful editing. The -Dsred lines were obtained by crossing to line PBac-transposase (BL32070) and the removal of the DsRed cassette was confirmed by sequencing.

#### Design of UAS-optoDronc without GFP

The p3xUAS-DRONC-linker-CRY2PHR was generated by amplifying DRONC-linker-CRY2PHR from UAS-optoDronc::GFP plasmid, generated in ^35^, by PCR and inserted in pJFRC4-3XUAS-IVS-mCD8::GFP (Addgene 26217) cut by NotI and XbaI (to excise mCD8::GFP) using NEBuilder HiFi DNA Assembly Method to generate p3xUAS-DRONC-linker-CRY2PHR. The construct was checked by sequencing and inserted at the attp site attp40A after injection by Bestgene.

The following primers were used for amplification:

*DRONC-Linker-CRY2PHR F:*

TAACCCTAATTCTTATCCTTTACTTCAGGCGGCCGCAACATGCAGCCGCCGGAGCTCGA

*DRONC-Linker-CRY2PHR R:*

CCACAGAAGTAAGGTTCC

### Live imaging and movie preparation

Notum live imaging was performed as followed: the pupae were collected at the fluid stage (0-10 hours after pupal formation), aged at 22°C or 29°C, glued on double-sided tape on a slide and surrounded by two home-made steel spacers (thickness: 0.64 mm, width: 20x20mm). The pupal case was opened up to the abdomen using forceps and mounted with a 20x40mm #1.5 coverslip buttered with halocarbon oil 10S. The coverslip was then attached to spacers and the slide with two pieces of tape. Pupae were collected 48 after clone induction and dissected usually at 16 to 18 hours APF (after pupal formation). The time of imaging for each experiment is provided in the table above. Pupae were dissected and imaged on a confocal spinning disc microscope (Gataca systems) with a 40X oil objective (Nikon plan fluor, N.A. 1.30), a Zeiss LSM880 (Zen Black software) equipped with a fast Airyscan using an oil 40X objective (N.A. 1.3), or a Zeiss LSM980 (Zen Blue software) equipped with a fast Airyscan 2 using an oil 40X objective (N.A. 1.3). Z-stacks (0.5 or 1 µm/slice) were captured every 1, 2, 5 or 10 min using autofocus at 22°C. The autofocus was performed using the plane of E-cad or other apical membrane marker as a reference, using a Zen Macro developed by Jan Ellenberg laboratory (MyPic) on the LSM880, the built-in autofocus function of Zen Blue on the LSM980, or a custom made Metamorph journal on the spinning disc. Movies were performed in the nota close to the scutellum region containing the midline and the aDC and pDC macrochaetae. Movies shown are adaptive local Z-projections. The plane of E-cad or other apical membrane marker was used as a reference to locate the surface of interest on spatial windows of the image (using the maximum in Z of average intensity or the maximum of the standard deviation). Other channels were then projected with a maximum intensity z-projection using an offset and projection stack size appropriate for the localisation of the signal of interest. This procedure was either applied using a custom Matlab routine^34, 81^. On some occasions where this local Z-projection pipeline could not accurately resolve folds in the tissue or would project the signal of the cuticle above the apical cell surface of interest, the Zellige^82^ Fiji plugin was used to locate and project the surface channel, and the resulting z-height maps were saved and imported into the custom local Z-projector Matlab routine as a reference surface from which other channels could be projected if necessary. Automation of the Zellige plugin to run on multiple timepoints was achieved with a custom ImageJ macro.

### Optogenetic control of caspase with optoDronc

Crosses involving expression of optoDronc were maintained in the dark at 25°C. Collection of pupae and dissection were performed on a binocular with LEDs covered by a home-made red filter (Lee colour filter set, primary red) after checking that blue light was effectively cut using a spectrometer.

#### Mild induction of optoDronc

For experiments involving mild induction of optoDronc, either globally, in regions of the tissue or in clones, blue light pulses were administered by acquiring images using the 488 nm laser either as a short time series (5 or 10 single planes within 5 minutes) or a single z-stack (10 planes of variable interval to cover the apical surface of the notum). optoDronc pulse images were acquired on a Zeiss LSM880 in fast airyscan imaging mode using an oil 40X objective with 488 nm laser power 2.8% and pixel dwell of 1.08µsec (for global activation of optoDronc) or 1.02µsec (for local and clonal activation).

#### Full activation of optoDronc

To assess the response of the effector caspase sensor GC3Ai to full and prolonged activation of caspase with optoDronc, z-stacks of 22 slices (1 µm/slice) were captured on a Zeiss LSM880 in fast airyscan mode every 2 mins using the 488 nm laser (2.8% power, pixel dwell 0.54µsec) to excite optoDronc and image GC3Ai intensity, and the 561 nm laser to image E-cad::mKate at the cell apical junctions.

### Image analysis

#### Cell segmentation

In order to extract quantitative data from individual cells over time, cells were segmented using the EPySeg Python package on the Z-projected E-cad surface. The model used for segmentation was either the built-in v2 model from EPySeg^39^, or a version of this model after re-training on our own previously corrected segmentation masks. Correction of segmentation masks was performed either using the Fiji plugin TissueAnalyzer, or the Napari plugin EpiCure^40^. Corrected segmentation masks were then exported for cell tracking.

#### Tracking using skimage and trackpy

For each of the large segmented GC3Ai movies, the input is an array of segmented images (I_k_)_k=1,…,N_, where k is a time-frame, and each image *I*_*k*_ is a snapshot of a tissue at time t = k × 5 min. Each image is a binary mask, where the value 1 (or 255) represents the boundary between two cells, and 0 is either intracellular space or background. The ‘skimage’ library of python was used to detect and label cells as connected regions in each image. Furthermore, the x,y position of each connected region in the image was saved as well as its geometric properties such as area, perimeter, eccentricity, solidity, orientation, and the coordinates of each pixel belonging to that connected region. All this was saved in one dataframe structure of the ‘pandas’ python library.

The ‘trackpy’ library of python was then used to track the connected regions over time. A pixel range of 15 was used with the adaptive step 0.95. Additionally, it was permitted for a cell to ‘disappear’ from the tracking during one time frame. This was done in order to compensate for some errors that occurred in the segmentation, such as when a real boundary between two cells was not detected in one image. The result was a dataframe as before, with a new label named ‘particle’, that maintains the same value for different time frames for the same connected region.

We eventually removed border cells, sensory organ precursor cells (SOP), and too short-lived cells from the analysis. Border cells were identified by pixels touching the edges of the image and were flagged as background. Then, using the ‘skimage’ library, the background regions were dilated with 3 x 3 kernels, and cells that overlapped with this dilated background were then flagged as border cells. Sensory organ precursor (SOP) cells are specialized mechanosensory cells that behave differently from their epidermal neighbours in the notum. Since SOPs grow to a larger size than their epidermal neighbours at late developmental stage, any cell that in any moment reached an area of 3000 pixels (89.79μm^2^) was removed from analysis. Any particle that lived less than 15 time frames (75 min) was also removed from the analysis.

Following this curation, the remaining tracked particles were those that lived long enough, were not too large (not SOPs) and were never at the edge of the tissue. These relevant cells were used for all further analysis.

#### Detecting dying cells using morphological features

The identification of dying cells was based on the identification of cells which do not appear in the final time frame, and which undergo apical constriction. The raw signal of apical area was smoothed to remove noise. As some tracked cells disappeared over one time frame, the ‘numpy’ library was used to interpolate the signal over those time frames which lack information about the cell. Then, a median filter of kernel size 15, from ‘scipy.signal’ library of python, was used to smooth the signal. Cells whose smoothed area shrank at least 50% over the last 75 min of their life were tagged as dying cells. Their raw area was then assessed to ensure that the area was in progressive decline in the last 4 time points, i.e. 20 min prior to disappearance.

#### Detection of area inflection point as the onset of cell extrusion

The onset of extrusion was defined as the moment when a dying cell starts reducing in size before its full delamination and disappearance from the tissue. A ‘two-line fit’ procedure was used to detect this moment automatically for each dying cell.

A given dying cell lives on a time interval [t_0_, t_end_], and its area is recorded on every 5-min time step in this interval. At times {t_0_, t_1_, t_2_, . . ., t_end_}, it has a recorded area, {a_0_, a_1_, a_2_, . . ., a_end_}. As the cell does not exist after *t*_*end*_, one more time step is defined: *t*_*end* + 1_ = *t*_*end*_ + 5 *min*, and the final size of the cell *a*_*end*+1_ = 0. The time observations were then split in two sets {t_0_, . . ., t_k_} and {t_k+1_, . . ., t_end+1_}, and using the python library ‘statsmodels.api’, two separate ordinary least squares models were fit on the time-area points. From both models the sum of all residuals was extracted, and then the residuals from the left-line and right-line fit were summed. This value was saved as *r*_*k*_.

This process was repeated for every k = 1, . . ., t_end_, producing a set of residuals {r_1_, . . ., r_end_}. At the minimum value in this set, *r*_*m*_, the corresponding *t*_*m*_ was defined as the inflection point of the cell area and the onset of extrusion for the given cell, *t*_*onset*_ = *t*_*m*_.

Some dying cells divide in the course of their life, and this method could wrongly attribute the moment of division as the onset of extrusion. For this reason, the inflection point analysis was restricted to the last hour (last 12 recorded points) of a dying cell life, so the time interval of interest was in fact [t_end_ − 60 min, t_end_].

#### Interpolation and smoothing of signals

It is worth noting that raw recordings of signals (area, perimeter, gfp levels) are noisy. Even though the quality of imaging is high, the raw signals will always have some amount of technical error. Parameters such as area, perimeter, eccentricity were smoothed using a median filter with kernel size 15, which was chosen preferentially over a mean filter due to the fact that it shortens the signal for the size of its kernel.

The GC3Ai signal (mean GFP intensity) was smoothed using a Savitzky-Golay filter, which fits successive subsets of adjacent data points with a low degree polynomial using the linear least squares. This was implemented using the ‘scipy.signal’ library of python with kernel size 15 and the 3^rd^ degree polynomial. Furthermore, such smoothing enables for an easy calculation of the derivative of the signal, in this case - the caspase levels. The derivative of the signal would be a derivative of the polynomial, and in practice, we only need to specify the parameter of the function.

While GC3Ai signal should cumulate in the cell, signal can also fluctuate due to adaptive z-projection which can generate some negative values in the GC3Ai derivative remaining close to 0. We decided to set all these values to 0.

After running the full image analysis pipeline on both movies separately and confirming that the same relationships and conclusions were drawn from each, the two datasets were pooled for greater statistical power and more information for machine learning after rescaling raw GC3Ai signal distribution using the median.

### Caspase threshold search and logistic regression

To investigate the existence of an overall threshold that determines if a cell in the tissue will extrude, all values of smoothed GC3Ai intensity and caspase activity from all cells, dying and non-dying, at all timepoints, were taken into account. This analysis was performed on more than 15 000 cells, where each cell lives at least 15 time frames and at most 192 time frames.

For a cell, *c*, we have values of a signal *x* = {*x*_*k*_}_*k*=*t*0_,..,*t*_*end*_, where *x*_*k*_ is a value of a signal (GC3Ai or caspase) at time frame k, *t*_0_ is a time frame when the cell *c* is first observed, and *t*_*end*_ is its last time frame.

Each *x*_*k*_ of every cell is assigned a value, labelled with

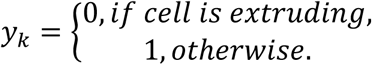

If a cell does not die during the movie, all values {*y*_*k*_}_*k*=*t*0_,..,*t*_*end*_ are equal to 0. If a cell dies during the movie, we identify its extrusion time *t*_*onset*_, then

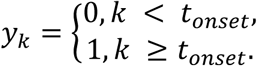

A logistic regression was performed on (x_k_, y_k_) pairs of values, fitting a logistic model (sigmoid function) to the data. This function takes real values and outputs a value between 0 and 1. We interpret it as probability to have a certain value of *x*_*k*_ to be labeled with 0 or 1. A good fit of the model and a sharp transition between the two states (not extruding and extruding) would indicate the existence of a threshold of the signal (either GC3Ai or caspase activity) that determines if the cell is extruding or not.

The goodness of fit of each logistic model to the real data was assessed by calculating the Tjur pseudo R-squared (Tjur’s R^2^, also known as the coefficient of discrimination). Tjur’s R^2^ compares the average fitted probability of the two outcomes, and is calculated as follows:

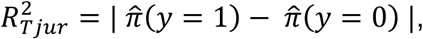

where 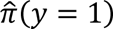 is the average predicted outcome from the model when y = 1, and 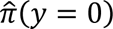 is the average predicted outcome when y = 0. The value of Tjur’s R^2^ can be interpreted as follows (see **Fig. S1** for more details):

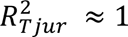: The model has absolute predictive power, values of the y variable are segregated perfectly by the x variable.

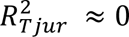: The model has no predictive power, values of the y variable are not segregated by the x variable.

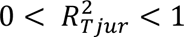: The model has some predictive power, values of the y variable are somewhat segregated by the x variable.

### Testing correlation between caspase activity at the onset of extrusion and GAL4 expression

To assess whether cell-to-cell variability in measured caspase activity at the point of commitment to death could actually be explained by differences in expression of the caspase sensor under the control of the GAL4-UAS system, both the UAS-GC3Ai caspase sensor and the constitutively expressed UAS-His3-IFP far-red histone marker were expressed in the same notum under the control of the notum-wide *pnr-G4* driver. Extrusion events were automatically detected using DeXtrusion^83^ and segmented using the TissueAnalyzer Fiji plugin^84^, and ROIs of segmented extruding cells were used to measure the evolution of perimeter and GC3Ai intensity over time. Cell perimeter was smoothed using a mean filter of kernel size 5 datapoints, and a custom MATLAB routine was used to automatically detect the perimeter inflection point using a two-line fit and assign this as the onset of extrusion (using the same logic as the approach detailed in methods section “Detection of area inflection point as the onset of cell extrusion”). Raw GC3Ai intensity of extruding cells was smoothed over 3 datapoints, the derivative of this signal was calculated to quantify the rate of change, and this derivative was smoothed over 3 datapoints (hereafter “caspase activity”). The cytoplasmic GC3Ai signal was used to follow the cell shape of extruding cells in z from their apical to nuclear planes and identify their respective His3-IFP-marked nuclei, maximum intensity local z-projections of which were segmented using the StarDist Fiji plugin^85^. Caspase at the onset of extrusion was plotted against His3-IFP mean intensity at the onset of extrusion, and the correlation was calculated using Spearman’s R.

### Correlation between cell parameters and caspase activity at the onset of extrusion

To assess the strength of the correlation between caspase activity at the onset of extrusion and other features of dying cells over time, the area inflection point of each dying cell was detected as previously described, and the cell’s value of caspase activity (smoothed derivative of mean GC3Ai intensity) at that timepoint was taken as its caspase activity at the onset of extrusion. All dying cells were aligned in time by their onset of extrusion (t^onset^), and their values of various morphological and caspase features in windows of time prior to the onset of extrusion were extracted. Specifically, for each dying cell, values of caspase activity, GC3Ai intensity, area, perimeter, derivative of area, derivative of perimeter and eccentricity were averaged in a 20-minute sliding window moved in 5-minute increments over the period covering -10hrs to 0hrs relative to the onset of extrusion. Using python, linear correlations were then calculated between the values of each parameter in each time window and caspase activity at the onset of extrusion, such that the correlation coefficient R could be computed for each relationship and used to populate a matrix showing the strength of the association between the level of active caspase at the onset of extrusion and the value of each parameter across time prior to extrusion.

### Mapping deaths following optoDronc activation

Coordinates of cell death events (x,y,t) following global mild optoDronc activation were registered manually by placing a point ROI at the coordinates of closure of an extruding cell in Fiji. The coordinates of the 4 anterior and posterior dorsocentral macrochaetae (aDC and pDC) in each notum were also manually registered to be used as landmarks for Procrustes transformation. The same process was carried out for control nota which expressed optoDronc but received no blue light pulses. Procrustes transformation was carried out using the built-in procrustes function in Matlab to take the landmark positions from each mapped notum, translate them to the same origin, scale them to the same size and rotate them until the coordinates of corresponding landmark positions aligned as closely as possible. This allowed the procrustes-transformed coordinates of extrusions from each notum to be superimposed on a common spatial map of the notum. The number of extrusions in discrete spatial windows was calculated by sorting points into bins by their Procrustes-transformed coordinates on a 15x15 density heatmap (covering roughly 5x5cells). A single density heatmap was generated summing all values for each group of nota. In the case of the density histograms shown in **Fig 1G**, the control map was then subtracted from the blue light pulses map through simple matrix subtraction to generate a map showing the spatial distribution of additional optoDronc-induced deaths.

To compare the spatial distribution of optoDronc-induced deaths at different developmental stages with the pattern of GC3Ai at the relevant stage, single images of GC3Ai distribution from the same notum at early and late stage of development were used to generate maps by averaging the fluorescent intensity of the sensor in spatial windows on a 15x15 grid (each box in the grid has width, height = 23.45µm). Maps of optoDronc-induced deaths from early and late pupae were generated as described above, this time using Procrustes transformation to align the coordinates of extrusions to the notum from which the GC3Ai map was generated as a reference. Furthermore, the limits of the extrusion density map were this time restricted to the limits of the GC3Ai reference image, to ensure a more like-for-like comparison of spatial windows between the two maps. Simple pairwise correlation between bins on the heatmaps of GC3Ai and extrusions at the relevant developmental stage was then calculated using Matlab to assess the strength of the relationship between the two classes of spatial map.

The extent to which optoDronc-induced deaths were clustered in the case of early, late and *hid* RNAi nota was assessed by measuring spatial autocorrelation of the summed maps of extrusion events from all nota in each group. Spatial autocorrelation was measured by calculating global Moran’s *I* using the moran.test function as implemented in the ‘spdep’ R package^86^. Global Moran’s *I* reports the extent to which the value of each position in the grid is related to the value of the neighbouring positions (here considering ‘rook’ neighbours, i.e. adjacent neighbours but diagonal ones), and therefore provides an index of clustering/dispersion. The value can be interpreted as follows:

0 < *I* < 1: data are more spatially clustered than expected from a random distribution

*I* ≈ 1: data are randomly spatially distributed

-1 < I < 0: data are more spatially dispersed than expected from a random distribution

### Calculating death probability following optoDronc pulse as a function of GC3Ai at t0

To assess the probability of cells in regions of high endogenous caspase to undergo cell death following mild optoDronc activation as a function of their cumulative caspase activity at t0, the GC3Ai sensor was combined with a non-GFP-tagged version of optoDronc. A single z-stack was captured at t0 to image E-cad::mKate and GC3Ai, and simultaneously trigger mild optoDronc activation (see methods section ‘mild induction of optoDronc’ for precise conditions of optogenetic activation). Cell contour was then imaged and projected over the following 90 minutes, and extrusions were manually tracked by placing a point ROI in the centre of each dying cell at the start of the movie. The image of E-cad::mKate and GC3Ai at t0 was segmented using EPySeg and corrected using the Fiji plugin TissueAnalyzer. The segmentation mask was then exported as a set of Fiji ROIs which were used to measure the mean intensity of GC3Ai within each cell. Extrusion probabilities were assigned by taking the coordinates of each manually tracked extrusion and measuring the Euclidean distance between this point and the centroid of each cell in the mask. The cell with the minimum distance between its centroid and this point was considered to be the dying cell, and its extrusion probability was set to 1, and after this was done for all extrusions, all other cells were assigned extrusion probability 0. Normalised GC3Ai values were calculated by dividing all GC3Ai intensity measurements by the maximum GC3Ai value from their respective movie, to allow data from multiple movies to be pooled. Values of normalised GC3Ai intensity were binned and the mean extrusion probability at each bin was plotted to show the relationship between cumulative caspase activity at the onset of optoDronc activation and probability of death in the following 90 minutes.

### Triggering past caspase activity in clones and calculating rate of death

To assess the difference in rate of death between WT clones and clones with past caspase activity within the same tissue, optoDronc was expressed in clones. Virgin female flies with the genotype *hsflp; E-cad::mKate/CyO; act>cd2>GAL4,UAS-cd4-mIFP* were crossed with males of the genotype *UAS-optoDroncGFP/CyO; MKRS/TM6B*. Clones were induced through a 15 minute heat shock in a 37°C water bath, and tubes were then maintained in the dark at 25°C. Fluid stage pupae were collected around 32 hours after clone induction and placed at 22°C, then imaged at 48 hours after clone induction. Mild optoDronc activation in clones was induced by capturing a single z-stack exposing the notum to blue light (see methods section ‘mild induction of optoDronc’ for precise conditions of activation), then the cell contour and clones were imaged for the following 5 hours using the E-cad::mKate and mcd4-IFP markers, respectively. All extrusions in the period of 1.5hrs – 5hrs following optoDronc activation were tracked manually by detecting the closure of delaminating cells, and each event was scored as either a WT or clonal death. We focused on the posterior region were caspase gets developmentaly activated later (see **movie S1**). The image of cell contour at 1.5hrs following optoDronc pulses was segmented using the python package EPySeg, and corrected using the napari plugin EpiCure. Cells at the border of the image were removed from the segmented cell mask, the number of clone cells was detected manually by counting cells in the mask expressing the mcd4-IFP marker, and the number of WT cells was calculating by subtracting the number of clone cells from the total number of cells in the mask. Data from two movies was pooled, and death rate was calculated for both WT and clonal populations by dividing the number of deaths in each population by cell number at 1.5hrs, then multiplying by 100 to be expressed as a percentage.

### Assessing rate of protein turnover with Proteoflux

To investigate any difference in the rate of protein turnover in distinct regions of the pupal notum, a pulse-chase experiment was carried out with the Proteoflux tool. Fluid-stage pupae expressing the Proteoflux tool under the control of a heat-shock promoter were selected, aged overnight at 22°C, then mounted on double-sided tape on a slide and expression of Proteoflux was induced by placing the slide in an incubator at 37°C for either 60 or 90 minutes. The pupae were then removed from the incubator, immediately dissected and imaged on a confocal spinning disc microscope (Gataca systems) with a 40X oil objective (Nikon plan fluor, N.A. 1.30) to capture the decay of the fluorescent signal.

### Machine learning analysis

#### Defining cell groups

To define cell groups for death prediction analysis, all cells touching a dying cell at its onset of extrusion (t^onset^) were designated as neighbours, and the collection of neighbour and dying cell was designated as a group. With 1,130 dying cells in the segmented dataset, as many groups were defined. The groups contain between 5 and 11 neighbours.

To define groups of neighbours, for a dying cell *c*^*D*^:

1. Find the time of the onset of extrusion, *t*^*D*^*_onset_*
2. From the time frame at *t*^*D*^*_onset_*

I. Get coordinates of all pixels of the dying cell *c*^*D*^, and create a mask *c*^*D*^, such that *c*^*D*^ is a matrix of the full image size, where each element of a matrix corresponds to a given pixel in the image and *c*^*D*^*_ij_* = 1 if a pixel (i, j) belongs to *c*^*D*^, and 0 otherwise.
II. Dilate the mask *c*^*D*^and find all cells in the same time frame that have some intersection with *c*^*D*^.
III. Verify if the intersecting cells are in the set of relevant cells, if yes - add them to the list of neighbours of *c*^*D*^.

#### Predicting dying cell using instant GC3Ai and caspase levels

Among groups containing a dying cell and all of its neighbours, we compared absolute GC3Ai intensity and caspase activity at the onset of the first extrusion and counted the proportion of groups where the first dying cell have the highest relative GC3Ai or caspase activity.

#### Time series forest (TSF)

The question of predicting which cell in the group of cells will be the first to die can be formulated as a time series classification problem, i.e. a machine learning problem. Due to the temporal structure of the input data, standard machine learning algorithms that are designed for structured data are usually not well suited. In particular, the order of the values is an essential part of time series, and consecutive time points are likely to be highly correlated.

Time series forest^43^ is a relatively intuitive algorithm. It derives summary features for all time series by dividing them into intervals and summarising each interval by its mean, standard deviation and gradient. Then a Random forest-like strategy is employed to select between a random subset of these features at each node in each of an ensemble of trees. A novel selection criterion is used that considers both entropy gain and the margin by which a feature separates the classes. This continues until the entropy gain ceases to improve, at which stage the node is defined as a leaf. TSF has been shown to be accurate classifier while being computationally efficient.

#### Training and testing data set

We want to investigate if the past behaviour of some cell variable could be informative in predicting which cell in a given group of cells is going to die first.

The groups of neighbours are defined as above - one dying cell and its neighbours at the time of onset. Now we look at the past behaviour of these cells. We have opted to use sliding-window strategy to investigate if and which signal is predictive of a dying cell. We define a time interval I_w,z_, with window size, w and ending position with respect to the onset of extrusion of a dying cell of a group, *t*_*onset*_ − *z*, so that

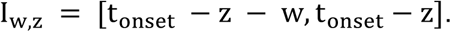

For example, for a time window of 1 hour that start 90 min prior to the onset of extrusion and ends 30 min prior to the onset, the interval we will look at is I_w,z_ = [t_onset_ − 90 min, t_onset_ − 30 min]. Now, if the dying cell of a group entered extrusion at, for example *t*_*onset*_ = 100 *min*, the time interval of interest for us will be *I*_*w*,*z*_ = [10 *min*, 70 *min*], i.e. in time frames [2, 14]. In absolute terms, each group of neighbours is analyzed based its own time of onset, so the actual time interval is group-specific, so the analysis is given in terms of ‘how far are we from the onset’.

For training TSF models we use only the information from one signal on the given interval. The algorithm to train one TSF model for given parameters w and z is as follows:

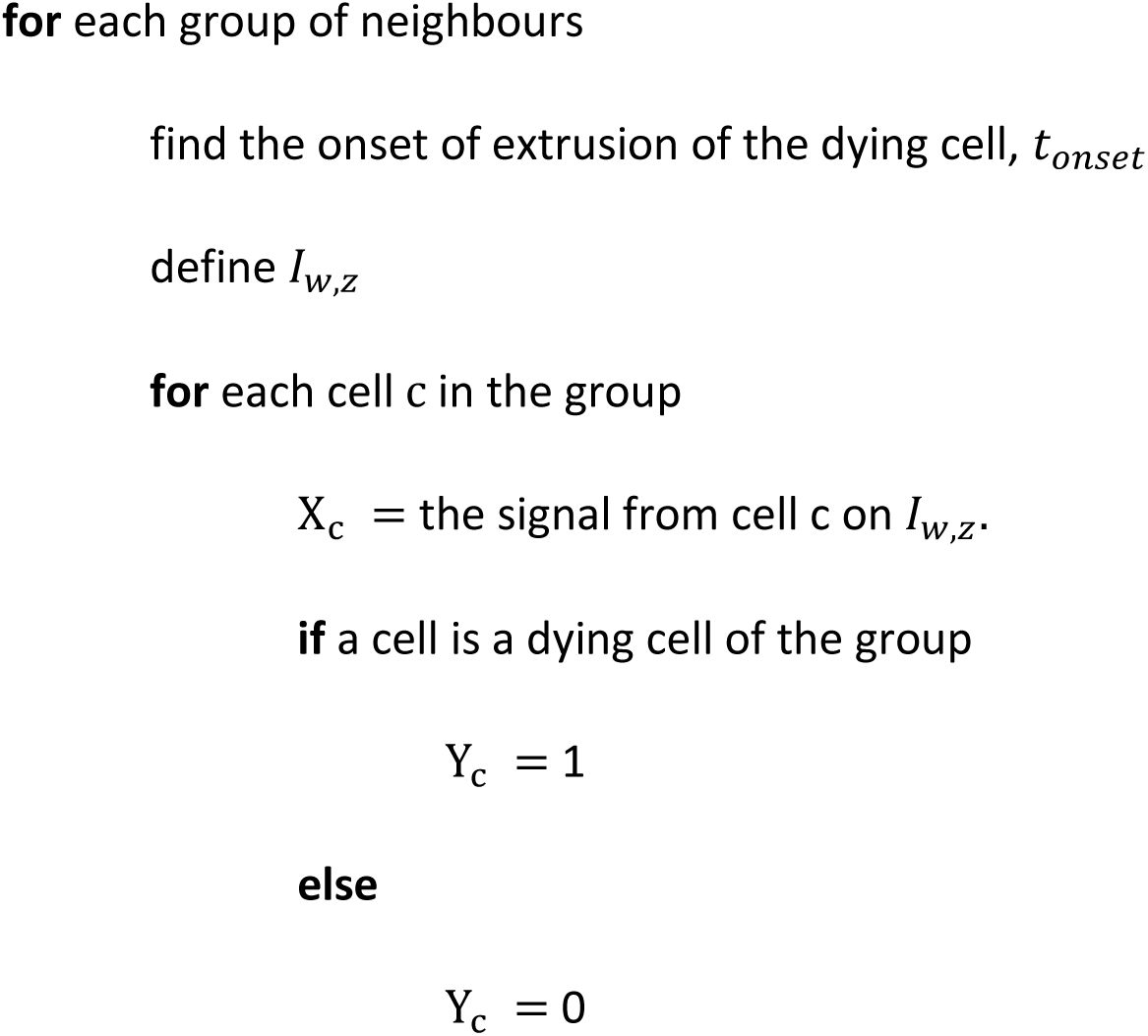

The full dataset is a list of all time series, X = {X_c_}, for all cells from all groups of neighbours, with corresponding classifications Y = {Y_c_}. If no other transformation is made, the neighbouring cells are related to each other through the definition of the time interval used for the extraction of the signal. Note that for every new set of parameters w and z we train different TSF model.

We use 70% of randomly picked data for training and the remaining 30% of data for testing. This means that if a given dying cell is selected to be used for training, all of its neighbours will be used for training.

#### Hyperparameters

We use the implemented version of TSF algorithm in python from **pyts** library [4]. There are a few hyper-parameters that need to be set for good model fitting. Most notably we need to set **n_estimators,** the number of trees in the forest. Following an example given in^87^, we use the *out-of-bag* error in order to estimate a suitable value for n_estimators at which the error stabilises. Finally, we use n_estimators = 125.

#### Training

We firstly trained TSF models using all neighbours in training datasets. For each TSF^w,z^ model, that is defined by the window size w and the final time point prior to onset of extrusion z, we train models based on the:

- GC3Ai signal, caspase activity, area, perimeter and eccentricity.

We vary parameters w and z as follows:

- w ∈ [1 hr, 2 hr, 3 hr, 4 hr],

- z ∈ [10 min, 20 min, 30 min, 1 hr, 2 hr, 3 hr, 4 hr].

Note that for larger values of parameters w and z we have less cells in datasets.

#### Balanced datasets

In a movie 1 we detected 514 dying cells. In a movie 2 we detected 616 dying cells. Each of these dying cells has between 5 and 11 neighbours at the onset of extrusion. For an easier interpretation we can use balanced datasets - for one dying, take only one non-dying neighbour. For balanced datasets we add one layer of randomisation of our dataset as we choose one random neighbour from each group of neighbours.

#### Scores

For testing how well the trained model performs on a new, never-seen, dataset we use previously selected test dataset. Then we ask a model to predict if the given cell in a test dataset is dying or living, i.e. the model is classifying the given time series into two classes, 0 or 1. The success can be measured with different scores. Firstly we can count how many times the model predicted positive outcome (class 1, i.e. dying cell) correctly - true positives TP, or wrongly - false positives FP. Then we count how many times the model predicted negative outcome (class 0, i.e. non-dying cell) correctly - true negatives TN, or wrongly - false negatives FN. Then the scores are defined as follows:

1. Accuracy = 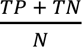, tells us how many true predictions have been made overall.
2. Precision = 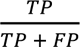, tells us how many true predictions have been made among all positive predictions.
3. Recall = 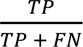, tells us how many true predictions have been made among all cases where the prediction should have been positive.
4. F1 score = 2 × precision × recall / (precision + recall), tells us how effectively a model makes a trade-off between precision and recall. Notice that F1 score goes to zero if any of the components goes to zero.

#### TSF prediction in a group of neighbouring cells using relative values

Another way to test and score a given TSF model is to ask it to predict the dying cell among a given group of cells. Rather than the overall classification of the time series, we are in fact interested to know which of the given cells (typically neighbouring cells) is the most likely to die first. Our metric will take a group of cells and instead of asking a model which class each cell belongs to, we ask the model which cell has the highest probability to be a dying cell.

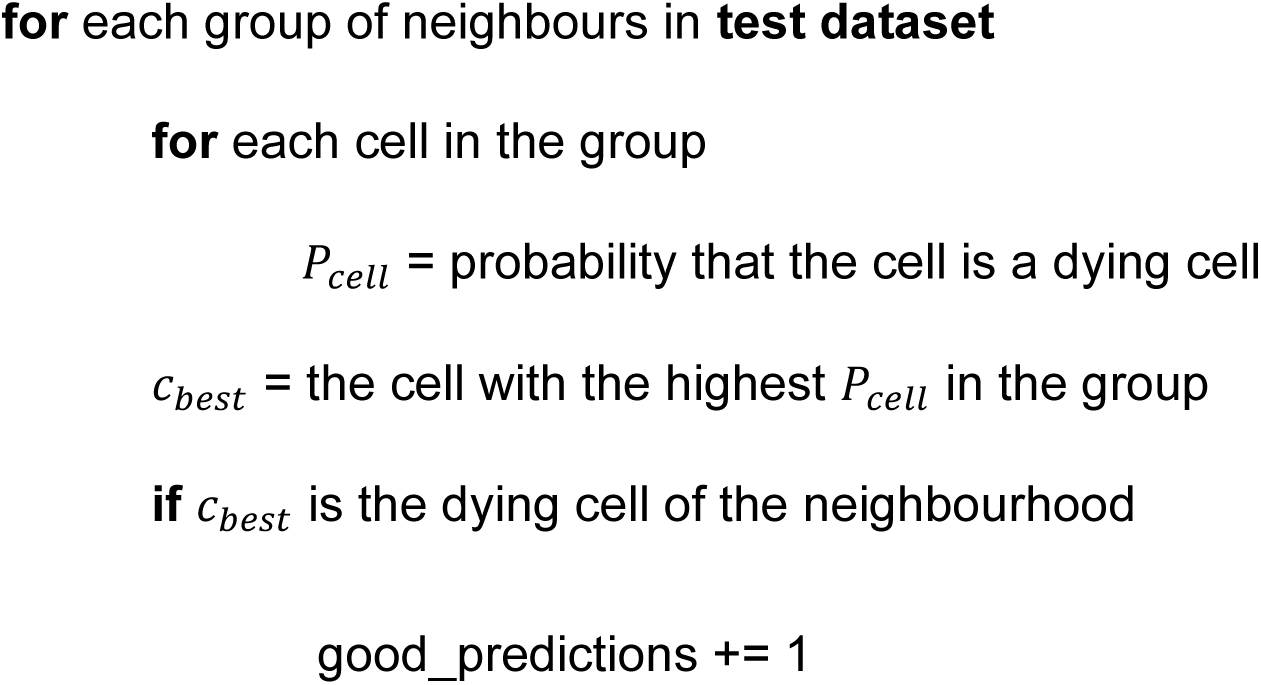

The maximum value for good_predictions is the total number of groups we have in the test dataset. The score is then defined as

local accuracy = good_predictions / total number of groups in dataset.

For this task, we used relative GC3Ai and caspase activity values between neighbours for training the TSF model. The modified datasets are created as follows,

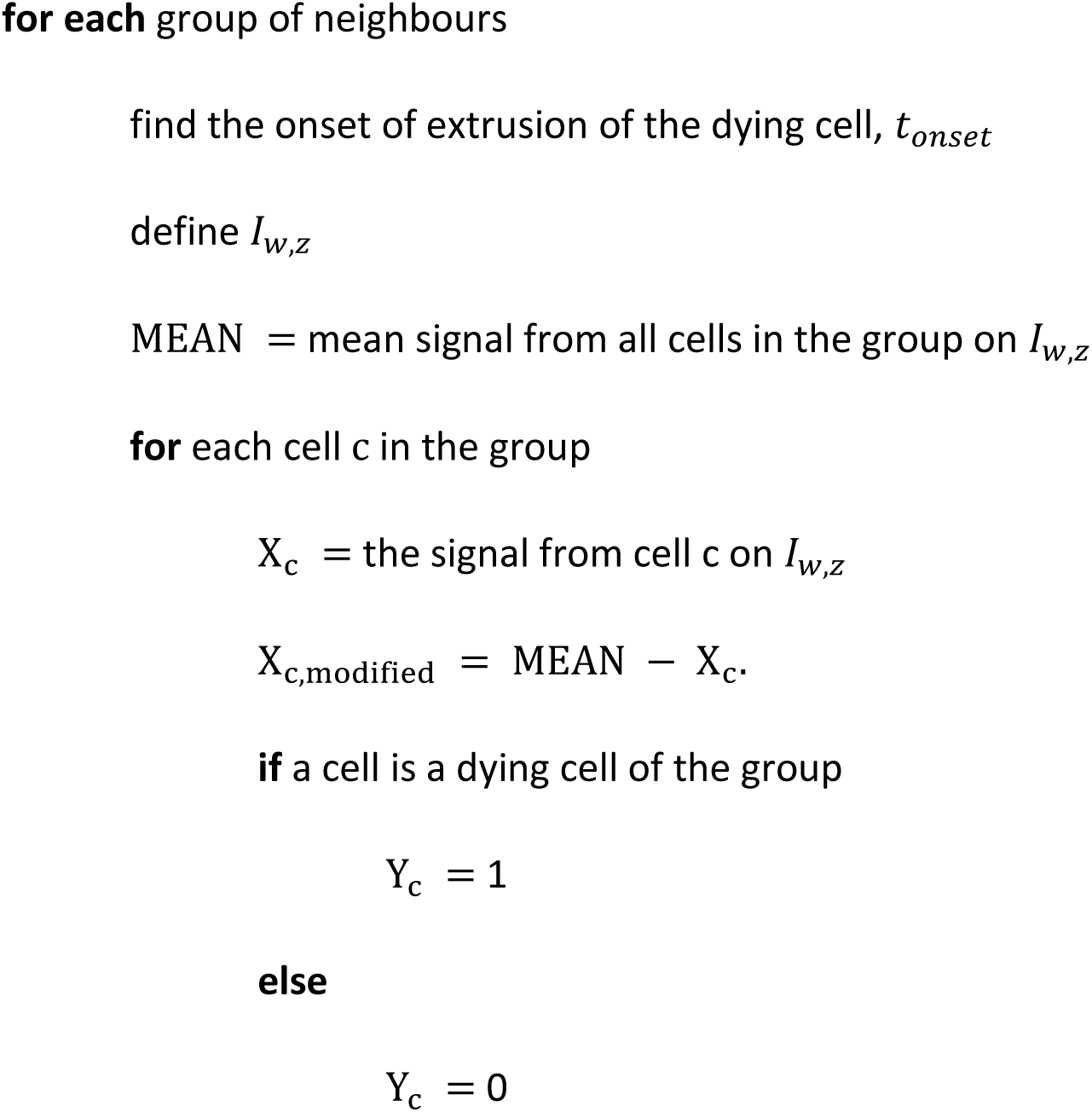

Then the procedure is the same as described above. We train TSF models on balanced datasets, using various signals, using w = 1hr and various values of parameter z. We calculate the scores and do local prediction on the test datasets.

### Statistics

Data were not analysed blindly. No specific method was used to predetermine the number of samples. The definition of n and the number of samples is given in each figure legend and in the table of the Experimental model section. Error bars are standard error of the mean (s.e.m.) or 95% confidence interval. p-values are calculated through t-test if the data passed normality test (Shapiro-Wilk test), or Mann-Whitney test/Rank sum test if the distribution was not normal. Statistical tests were performed on Graphpad Prism 8 or Matlab.

---

**Supplementary Figure 1:**
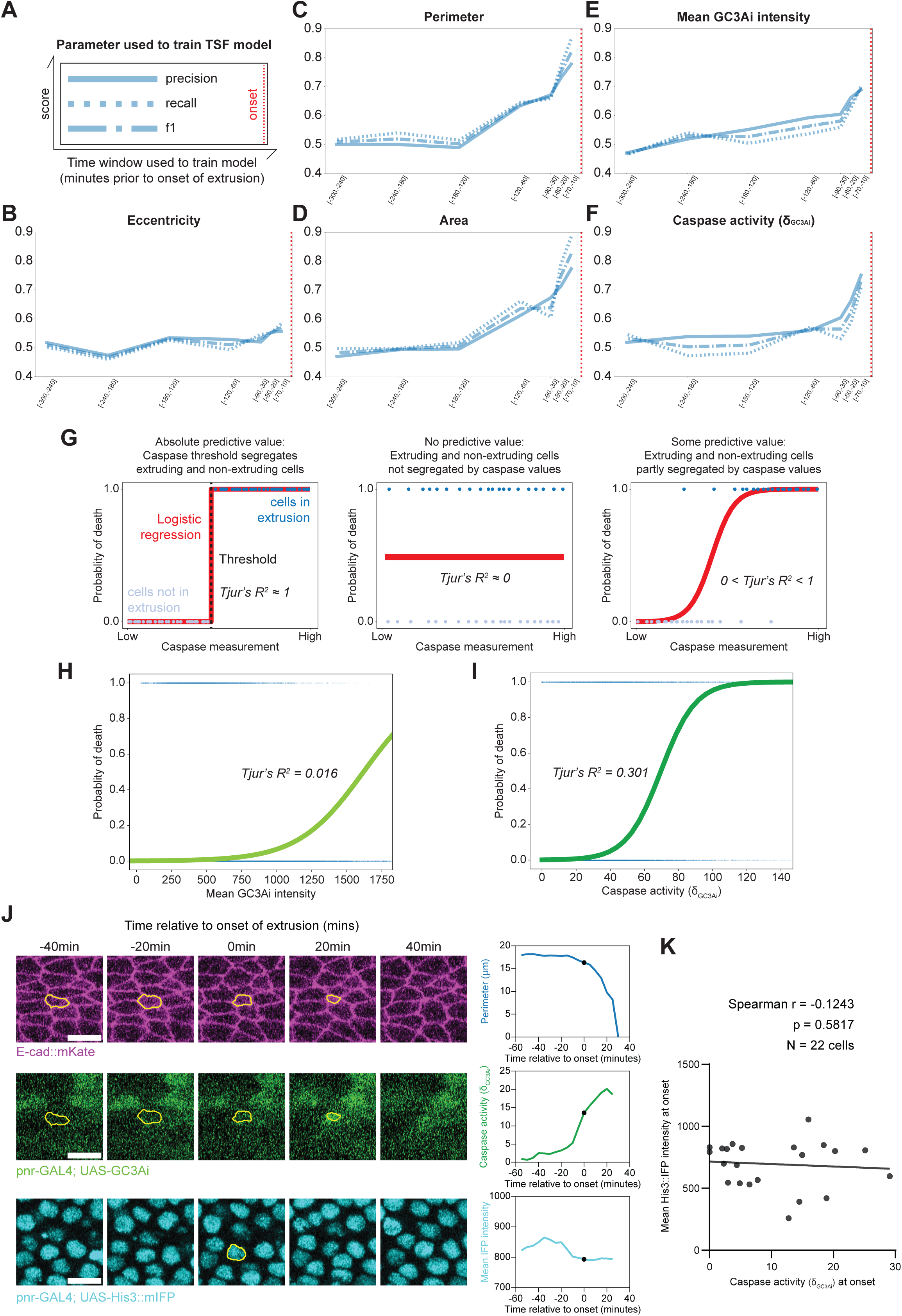
Identification of parameters and time windows relevant for cell death prediction (associated with Figure 1). **A:** Summary of the metrics shown in each graph of the Time Series Forest (TSF) prediction. Solid curve, precision, dotted curve, recall, dotted-dash curve, F1 score on the y axis. x axis shows the time windows used to train the model (in minutes prior to extrusion onset, red dotted line). **B-F:** Precision, recall and F1 score of the prediction of dying cells in the pupal notum using TSF models trained on the corresponding time window (x-axis) using a dying cell and one of its neighbours as control measuring cell eccentricity (**B**), cell apical perimeter (**C**), cell apical area (**D**), GC3Ai raw intensity (**E**) and caspase activity (GC3Ai intensity derivative, **F**). The baseline (random prediction) is at 0.5. **G:** Schematic of the segregation of data based on logistic regression and sorting of the dying and non-dying cells based on caspase activity. Left shows a perfect prediction (step-like function), middle the absence of predictive value, and right a partial predictive value. **H,I:** Logistic regression of the probability of cell death as a function of raw GC3Ai intensity **(H)** and caspase activity **(I)** based on the curves shown in Fig. 1 **D** and **E**. The discrimination power is estimated using Tjur’s R^2^. **J:** Snapshots of a single cell undergoing cell extrusion showing E-cad::mKate (top, magenta), UAS-GC3Ai signal (middle, green) and UAS-His3::mIFP (bottom, cyan) both driven by the pnr-Gal4 driver. Time 0 is the onset of cell extrusion. Scale bars=10μm. Blue curve shows the perimeter over time, green curve the caspase activity (GC3Ai derivative) and cyan curve the His3::mIFP intensity. **K:** Correlation between caspase activity at the onset of extrusion (x-axis) and His3::mIFP intensity (y-axis). n=22 extruding cells (one dot=one extruding cell). Note the absence of correlation between the far-red signal (a proxy of Gal4 transcriptional activity) and the caspase activity at the onset, r=-0.12, p=0.58.

**Supplementary Figure 2:**
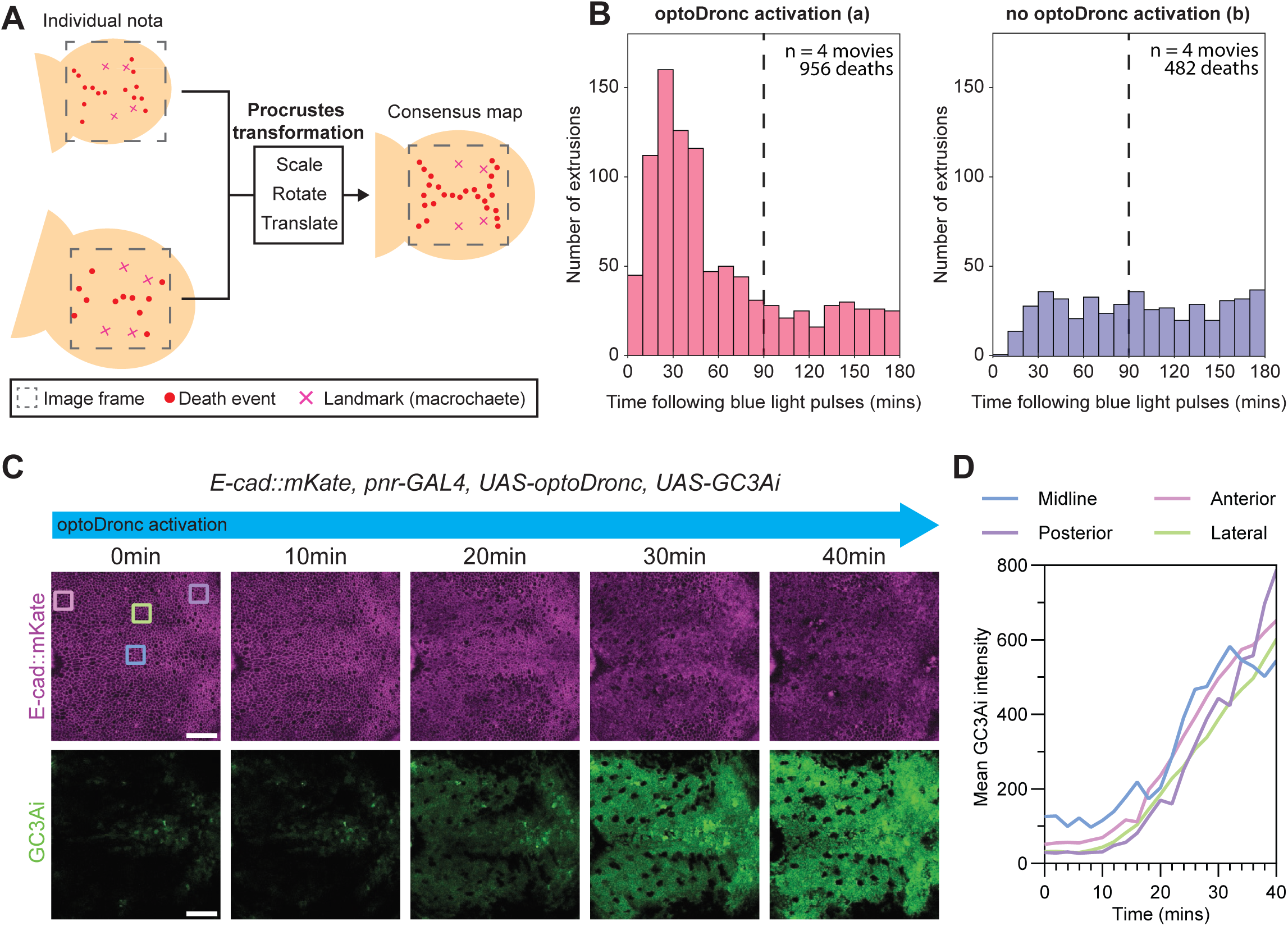
Susceptibility to caspase is spatially patterned (associated with Figure 1 and Figure 2). **A:** Schematic of the process used to align pupae upon optoDronc pulses to produce a consensus spatial map of optoDronc-induced deaths. Procrustes transformation is used to align, scale and rotate pupae and their corresponding mapped extrusions using 4 posterior macrochaetae (aDC and pDC macrochaetae) as spatial landmarks (pink crosses). **B:** Bar plots of the absolute number of cell extrusions counted at various time windows (minutes post-blue light activation) in *pnr-Gal4 UAS-optoDronc* pupae following mild and global activation of optoDronc (left, (a)) or in absence of blue light (right, (b)) (associated with Fig. 1F**,G**). n=4 pupae, 956 cell death after blue light activation, 482 in the controls. Note that the additional deaths observed in the blue light-activated pupae mostly occur in the first 90 minutes (black doted lines) and death rate returns to control levels after 90 minutes. **C:** Snapshots of a pupal notum expressing *UAS-optoDronc* (no GFP tag) and *UAS-GC3Ai* with *pnr-Gal4* upon global and persistent blue light exposure. E-cad::mKate in magenta, GC3Ai in green. Note the global and homogeneous increase of GC3Ai signal (despite the initial heterogeneity corresponding to the normal pattern of caspase activation. The coloured squares show the regions of intensity measurement quantified in **D**. Scale bars=50μm. See also **movie S4**. Anterior, left, posterior, right. **D:** Time evolution of GC3Ai intensity in different ROIs of the pupae shown in **C**, blue: midline, pink: anterior, purple: posterior, green: lateral.

**Supplementary Figure 3:**
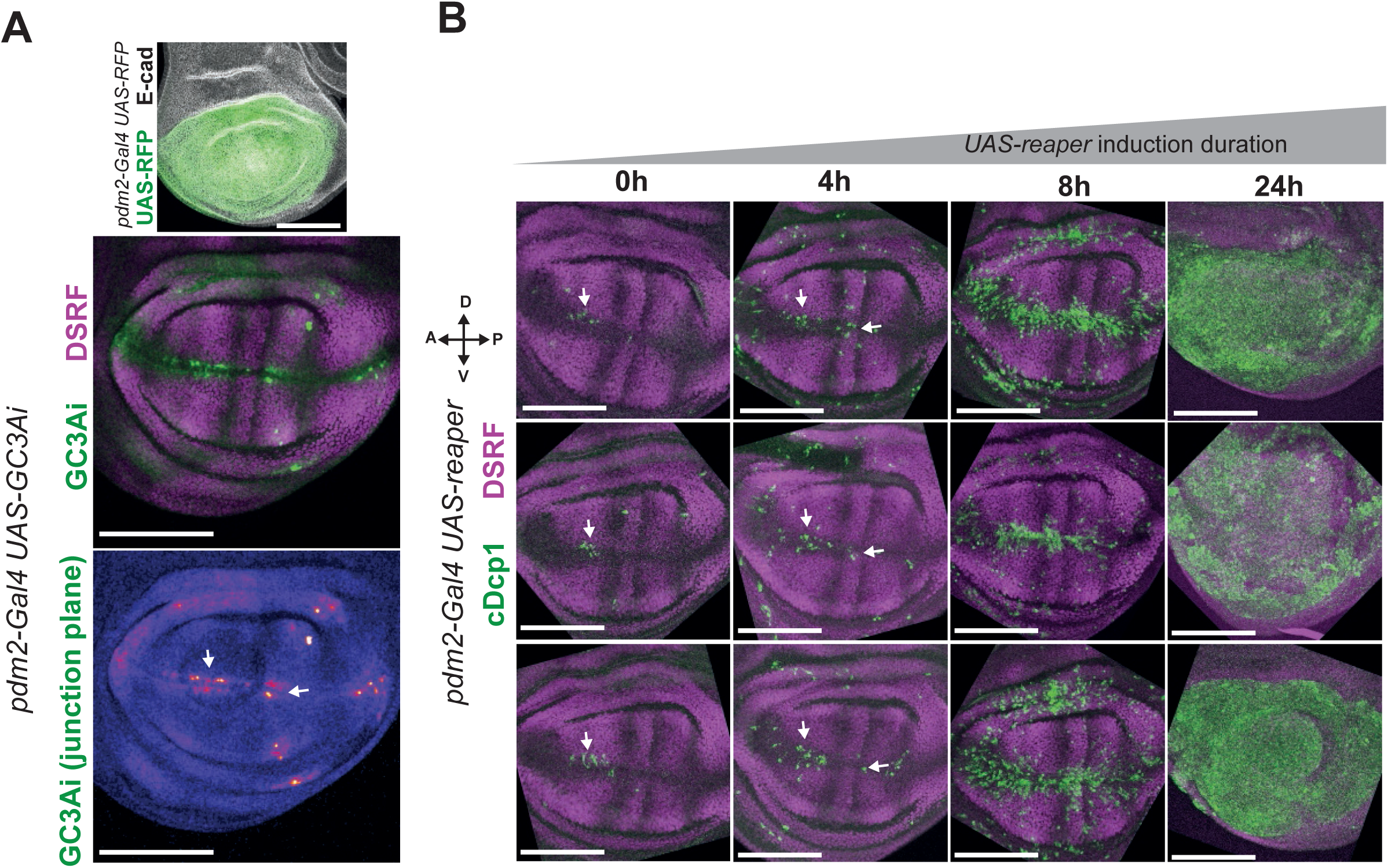
Susceptibility to caspase is also spatially patterned in the larval wing imaginal disc (associated with Discussion). **A:** Top, pattern of expression of the pdm2-Gal4 driver in the larval wing imaginal disc (green, UAS-RFP, grey, E-cad). Middle: local projection of DSRF staining (intervein regions, magenta), GC3Ai signal in the plane of adherens junctions (green). Bottom: Pseudo colour of GC3Ai intensity in the adherens junction plane. White arrows show regions of high GC3Ai signal. Scale bars=100μm. **B:** Representative examples of cell death distribution (cleaved Dcp1, green) upon induction of Reaper in the *pdm2-Gal4* domain for different activation time (duration of 29°C incubation in hours releasing Gal80^ts^ repression). Cell death is mostly located in the previously characterised apoptotic hotspots at 0h (see ^50^), is first enriched in regions already showing some cumulated GC3Ai signal after 4 and 8h of activation, and eventually is distributed throughout the wing disc. n=23, 24, 28 and 14 wing discs. Scale bars=100μm.

**Supplementary Figure 4:**
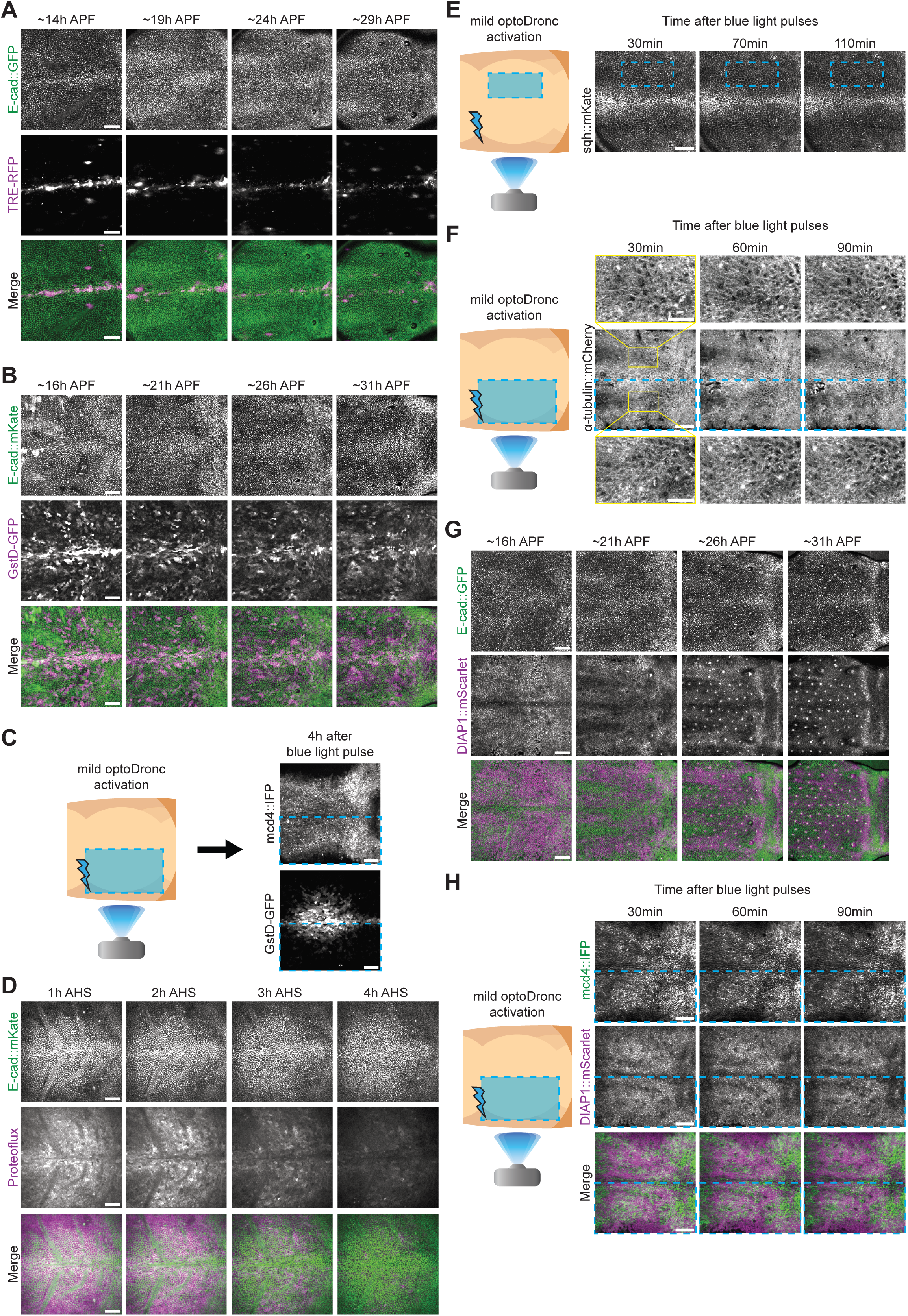
Testing putative regulators of the memory of past caspase activation (associated with Discussion). **A:** Snapshots of a pupal notum expressing the JNK reporter TRE-RFP (middle, magenta) and E-cad::GFP (top, green) at different developmental times (hours After Pupal Formation, APF). Anterior, left, posterior, right. Note the absence of signal in the posterior region. Scale bar=50μm. Representative of 2 pupae. **B:** Snapshots of a pupal notum expressing the ROS reporter GstD1-GFP (middle, magenta) and E-cad::GFP (top, green) at different developmental times (hours After Pupal Formation, APF). Anterior, left, posterior, right. Note the absence of clear signal upregulation in the posterior region. Scale bar=50μm. Representative of 3 pupae. **C:** Snapshots of a pupal notum expressing the ROS reporter GstD1-GFP (greyscale) and *pnr-Gal4 UAS-optoDronc* upon mild activation of caspase in the left region of the notum (see left schematic, doted blue rectangle) 4 hours after blue light activation (note that GstD1-GFP transcriptional reporter should perdure for hours upon activation). No clear difference can be observed between the activated and control regions. Anterior, left, posterior, right. Scale bar=50μm. Representative of 2 pupae. **D:** Snapshots of a pupal notum expressing the proteasome activity reporter (ProteoFLUX, hs-CL1::GFP, magenta) and E-cad::mKate (top, green) over time after heat shock induction of the proteasome reporter (hours After Heat Shock, AHS). Anterior, left, posterior, right. Note the absence of clear pattern of speed of intensity decrease (a proxy of proteasome activity), especially in the posterior region. Scale bar=50μm. Representative of 7 pupae. **E:** Snapshots of a pupal notum expressing sqh::mKate (greyscale) and *pnr-Gal4 UAS-optoDronc* upon mild activation of caspase in the right region of the notum (see left schematic, doted blue rectangle) over time after blue light activation (minutes). No clear differences of junctional sqh::mKate levels could be observed between the activated and control regions. Anterior, left, posterior, right. Scale bar=50μm. Representative of 5 pupae. **F:** Snapshots of a pupal notum expressing UAS-α-tub::mCherry (greyscale) and *pnr-Gal4 UAS-optoDronc* upon mild activation of caspase in the left region of the notum (see left schematic, doted blue rectangle) over time after blue light activation (minutes). Close up views of the two regions in yellow rectangles are shown above and below. No clear differences of cortical microtubule organisation could be observed between the activated and control regions. Anterior, left, posterior, right. Scale bar=50μm in main image and 20μm in cropped insets. Representative of 2 pupae. **G:** Snapshots of a pupal notum expressing the knock-in fusion of Diap1::mScarlett (magenta, middle) and E-cad::GFP (top, green) at different developmental times (hours After Pupal Formation, APF). Anterior, left, posterior, right. Note the absence of clear downregulation of Diap1 concentration in the posterior region where cell deaths occur from 16 to 26 hours APF. Scale bar=50μm. Representative of 2 pupae. **H:** Snapshots of a pupal notum expressing the knock-in of Diap1::mScarlett (middle, magenta), *pnr-Gal4 UAS-optoDronc* and UAS-mCd4::mIFP (top, green) upon mild activation of caspase in the left region of the notum (see left schematic, dotted blue rectangle) over time after blue light activation of caspases (minutes). No clear differences of Diap1 levels could be observed between the activated and control regions. Anterior, left, posterior, right. Scale bar=50μm. Representative of 2 pupae.

## Supplementary movie legends

**Movie S1:** Movie related to **Fig. 1A-E, 2A-C, 4A-C, S1A-I**. ∼16h APF notum local z-projection with E-cad::mKate (top left, magenta) and UAS-GC3Ai (top middle, green) driven by the pnr-Gal4 driver. Bottom left shows the skeletonised tissue. Bottom middle the cell tracking (1 colour per cell), the bottom right the colour coded cell average GC3Ai intensity (dark blue low, yellow high). Anterior, left, posterior, right. Scale bar is 50μm.

**Movie S2:** Movie related to **Fig. 1C**. Example of an extruding cell from a notum local z-projection with E-cad::mKate (middle, magenta) and UAS-GC3Ai (right, green) driven by the pnr-Gal4 driver. The skeletonised contour of cells is shown on the left, the dying cell is shown in white and with yellow contours. Anterior, left, posterior, right. Scale bar is 5μm.

**Movie S3:** Movie related to **Fig. 1F,J**. ∼20h APF notum local z-projection with E-cad::mKate (greyscale) expressing UAS-optoDronc::GFP driven by the pnr-Gal4 driver after global mild activation of optoDronc (left) or without blue light activation (control, right). Red dots show the dying cells. Anterior, left, posterior, right. Scale bars are 50μm.

**Movie S4:** Movie related to **Fig. S2C,D**. ∼20h APF notum local z-projection with E-cad::mKate (left, magenta) expressing UAS-optoDronc and UAS-GC3Ai (middle, green) driven by the pnr-Gal4 during a strong and global activation of optoDronc. Anterior, left, posterior, right. Scale bar is 50μm.

**Movie S5:** Movie related to **Fig. 2D**. ∼20h APF notum local z-projection with E-cad::mKate (greyscale) expressing UAS-optoDronc::GFP driven by the pnr-Gal4 after a local activation of optoDronc (in the red and blue square regions). The blue square was already exposed to a pulse of blue light 1 hour before. Dying cells in the regions of interest are shown with red dots. Anterior, left, posterior, right. Scale bar is 50μm.

**Movie S6:** Movie related to **Fig. 3A**. Nota local z-projection with E-cad::mKate (greyscale) expressing UAS-optoDronc::GFP driven by the pnr-Gal4 driver after global mild activation of optoDronc in a young pupae (left, 18h APF), late pupae (middle, 24h APF) or upon inhibition of *hid* (*UAS-hid* dsRNA, right). Red dots show the dying cells. Anterior, left, posterior, right. Scale bars are 50μm.

**Movie S7:** Movie related to **Fig. 3B**. ∼16h APF notum local z-projection with E-cad::mKate (left, magenta) expressing UAS-GC3Ai (middle, green) and UAS-hid dsRNA driven by the pnr-Gal4 driver. Compare with **Movie S1** for control. Anterior, left, posterior, right. Scale bar is 50μm.

**Movie S8:** Movie related to **Fig. 3D,E**. ∼16h APF notum local z-projection with E-cad::mKate (magenta) expressing UAS-GC3Ai (green) and UAS-optoDronc driven by the pnr-Gal4 driver after global mild activation of optoDronc (also used to image GC3Ai at T0). White dots show the dying cells. Anterior, left, posterior, right. Scale bar is 20μm.

**Movie S9:** Movie related to **Fig. 4D**. ∼24h APF notum local z-projection with E-cad::mKate (left, magenta), UAS-optoDronc::GFP and UAS-mCd4::mIFP-T2A-HO1 (green, middle) expressed in clones after mild activation of optoDronc. Red dots show the dying cells detected between 1.5hrs and 5hrs after the blue light pulse. Anterior, left, posterior, right. Scale bar is 30μm.

**Movie S10:** Movie related to **Fig. S4A**. ∼18h APF notum local z-projection with E-cad::mKate (left, green) and the JNK pathway sensor TRE-RFP (middle, magenta). Anterior, left, posterior, right. Scale bar is 50μm.

**Movie S11:** Movie related to **Fig. S4B**. ∼18h APF notum local z-projection with E-cad::mKate (left, green) and the ROS sensor GstD-GFP (middle, magenta). Anterior, left, posterior, right. Scale bar is 50μm.

**Movie S12:** Movie related to **Fig. S4D**. ∼16h APF notum local z-projection with E-cad::mKate (left, green) and the proteasome activity sensor Proteoflux (hs-CL1::GFP, middle, magenta) levels after a pulse of transcriptional activation. Anterior, left, posterior, right. Scale bar is 50μm.

**Movie S13:** Movie related to **Fig. S4D**. ∼20h APF notum local z-projection with sqh::mKate (greyscale) and UAS-optoDronc::GFP driven by the pnr-Gal4 driver after mild blue light stimulation in the blue square region. Anterior, left, posterior, right. Scale bar is 50μm.

**Movie S14:** Movie related to **Fig. S4F**. ∼20h APF notum local z-projection with UAS-αtub::mCherry (greyscale) and UAS-optoDronc::GFP driven by the pnr-Gal4 driver after mild blue light stimulation in the blue square region. Anterior, left, posterior, right. Scale bar is 50μm.

**Movie S15:** Movie related to **Fig. S4G**. ∼16h APF notum local z-projection with E-cad::GFP (green, left) and Diap1::mScarlet (KI) (magenta, middle). Anterior, left, posterior, right. Scale bar is 50μm.

**Movie S16:** Movie related to **Fig. S4H**. ∼20h APF notum local z-projection expressing UAS-optoDronc::GFP and UAS-mCd4::mIFP-T2A-HO1 (green, left) and Diap1::mScarlet (KI) (magenta, middle) with the pnr-Gal4 driver after mild activation of optoDronc in the blue square region. Anterior, left, posterior, right. Scale bar is 50μm.

